# Theoretical Tool Bridging Cell Polarities with Development of Robust Morphologies

**DOI:** 10.1101/328385

**Authors:** Silas Boye Nissen, Steven Rønhild, Ala Trusina, Kim Sneppen

## Abstract

Despite continual renewal and damages, a multicellular organism is able to maintain its complex morphology. How is this stability compatible with the complexity and diversity of living forms? Looking for answers at protein level may be limiting as diverging protein sequences can result in similar morphologies. Inspired by the progressive role of apical-basal and planar cell polarity in development, we propose that stability, complexity, and diversity are emergent properties in populations of proliferating polarized cells. We support our hypothesis by a theoretical approach, developed to effectively capture both types of polar cell adhesions. When applied to specific cases of development gastrulation and the origins of folds and tubes our theoretical tool suggests experimentally testable predictions pointing to the strength of polar adhesion, restricted directions of cell polarities, and the rate of cell proliferation to be major determinants of morphological diversity and stability.

## INTRODUCTION

Multicellular organisms are amazing in their ability to maintain complex morphology in face of continuous cell renewal and damages. Adult salamander can regenerate entire limbs (Eguchi et al. 2011), and, during development, some regions can maintain patterning when moved to different parts of an embryo or if the size is varied (Lyons, Kaltenbach, and McClay 2011). Given the vast complexity and diversity of living shapes, how can we reconcile the robustness to perturbations with flexibility to diversify? While undoubtedly the end result is encoded in the DNA and protein networks, looking for an answer at this level is challenging. Examples of phenotypic plasticity (Libby and Rainey 2011), convergent evolution, and contrasting rates of morphological and protein evolution (Cherry et al. 1979) show that morphological similarity may not couple to the protein sequence similarity (Stephen Jay Gould 1970). Inspired by the unfolding of morphological complexity in development, we propose that cellular polarity may be the key for reconciling complexity, robustness, and diversity of organismal morphologies.

During early development the increase in morphological complexity coincides with the progressive polarization of cells - first apical-basal (AB) polarity and then planar cell polarity (PCP) (Müller and Bossinger 2003; Roignot, Peng, and Mostov 2013; Andrew and Ewald 2010; R. Li and Bowerman 2010). This theme is ubiquitous across vertebrates and invertebrates (Figure 1): starting from the single fertilized egg cell, first the morula formed by non-polarized cells turns into the blastula - a hollow sphere of cells with AB polarity. Then, as cells acquire additional PCP, primary head-tail axis forms and elongates during gastrulation and neurulation (Loh, van Amerongen, and Nusse 2016). Because of the optical transparency these stages are particularly prominent in sea urchin. At the morula stage, a lumen in the center is formed and is gradually expanding as cells proliferate and rearrange into the hollow sphere. Next, during gastrulation, a group of cells invaginate and rearrange into a tube that narrows and elongates primarily by cell rearrangement and convergent extension movements (Martik and McClay 2017). The tube then merges with the sphere at the side opposite to invagination, and as a result, the sphere transforms into a torus. Emerging data suggest that PCP drives both invagination and tube elongation (Nishimura, Honda, and Takeichi 2012; Croce et al. 2006; Long et al. 2015) - a recurring theme across species.

**Figure 1.**
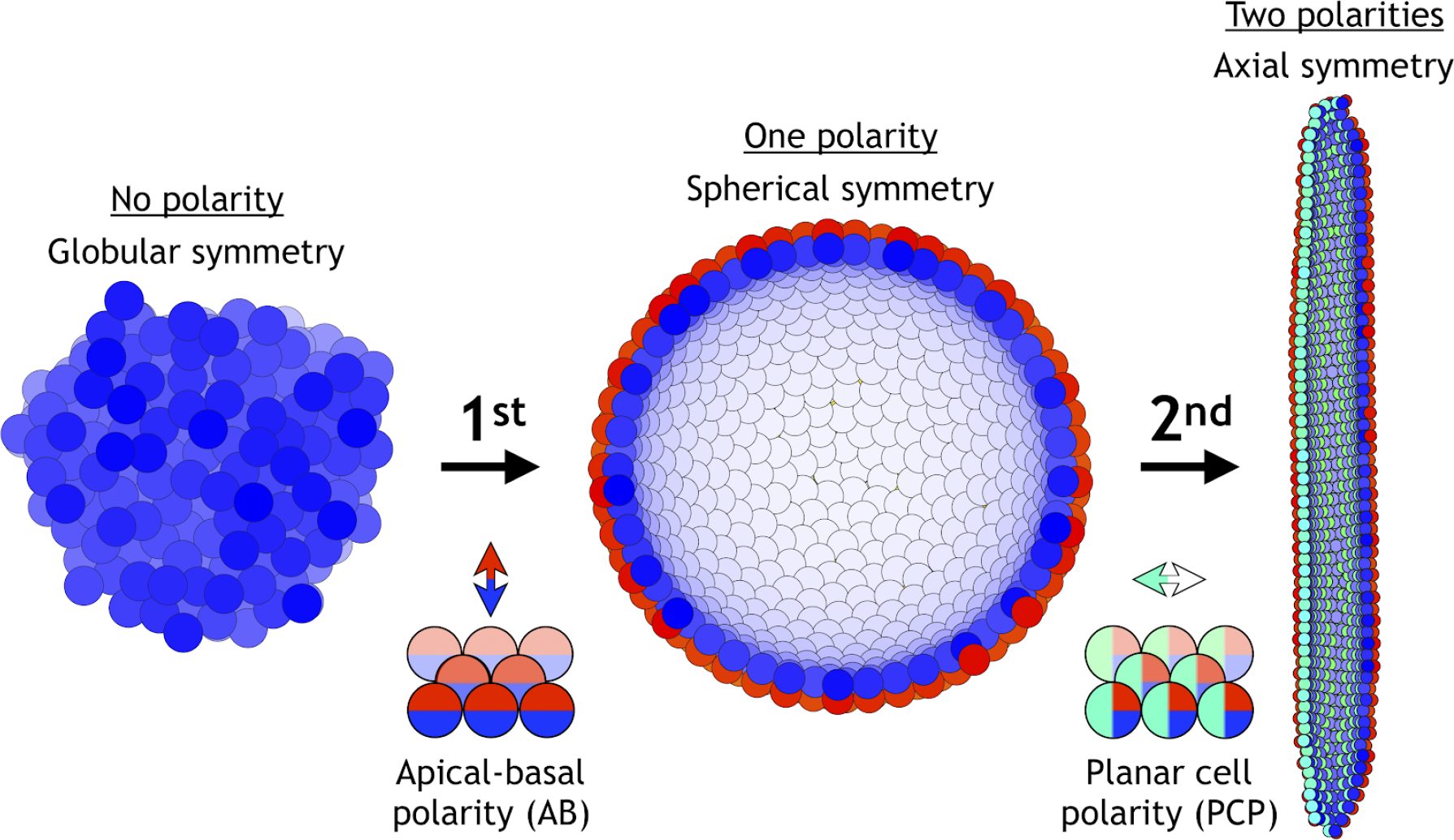
Two symmetry-breaking events, gain of apical-basal (AB) polarity and planar cell polarity (PCP), on cellular level coincide with the appearance of a rich set of morphologies. Starting from an aggregate of non-polarized cells (globular symmetry), individual cells can gain AB polarity and form one or multiple lumens (spherical symmetry). Additional, gain of PCP allows for tube formation (axial symmetry). Complex morphologies can be formed by combining cells with none, one, or two polarities. In Figure S1, we schematically illustrate how existing models capture different elements of development.

Mutations in PCP pathways produce shorter and wider tubes (Ochoa-Espinosa, Baer, and Affolter 2012; Saburi et al. 2008; Kunimoto et al. 2017), somites (Song et al. 2010), and embryos (Gong, Mo, and Fraser 2004). Formation and elongation of the tubes can proceed without cell division and cell death by cells rearranging along the tube’s axis, termed convergent extension (CE) (Andrew and Ewald 2010; Tanimizu, Miyajima, and Mostov 2009; Martik and McClay 2017). While the importance of PCP in gastrulation and tubulogenesis is well established (Andrew and Ewald 2010; Tanimizu, Miyajima, and Mostov 2009; Martik and McClay 2017; Kunimoto et al. 2017), it is unclear how polarity may control tube morphology.

The bulk-lumens-folds/tubes transition seen across animal species in early embryogenesis, is also a key feature of the later organ formation. The early stages of organogenesis in liver, kidney, brain, gut, and pancreas are apparently so robust, that they can be recapitulated *in vitro*, allowing for advanced quantification and manipulation (Little 2017). The case of pancreatic organoids is interesting as it illustrates an increase of morphological complexity from spheres to folds. Cells in *in vitro* pancreatic organoids first grow as a bulk and later acquire AB polarity and develop lumens. Depending on the growth conditions organoids develop into a hollow sphere or acquire a complex folded shape (Greggio et al. 2013). It is currently unknown what drives the transition from sphere to folded state; the two possible hypotheses are rapid proliferation or physical pressure by growing into a stiff matrigel.

Is the apparent link between cellular polarity and morphological complexity accidental? Or, could it be that morphological transitions, stability, and diversity are emergent features in a population of proliferating polarized cells? If true, can we identify what drives the transition from lumens to folds and tubes? Why are these stable? Can we predict what controls fold depth, and tube length and width? To answer these questions, we lack a unified approach that could bridge polar interactions between single cells to the global features emerging on the scale of thousands of cells in 3D.

Starting with D’Arcy Thompson’s seminal contribution (Thompson and Others 1942), quantitative models aided in understanding specific morphogenetic events (Figure S1). Among these are *invagination* (Odell et al. 1981; Rauzi et al. 2015; Polyakov et al. 2014; Hočevar Brezavšček et al. 2012), *primitive streak formation* (Newman 2008), *convergent extension* (Collinet et al. 2015; Belmonte, Swat, and Glazier 2016), *epithelial folding* (Buske et al. 2012; Osterfield et al. 2013; Monier et al. 2015; Murisic et al. 2015), emergence of global PCP alignment from local cell-cell coupling (Amonlirdviman et al. 2005; Le Garrec, Lopez, and Kerszberg 2006; Burak and Shraiman 2009), origins of *tubulogenesis* (Engelberg et al. 2008), and recently statistical properties of branching morphogenesis (Hannezo et al. 2017). However, they are often on either of the two ends of the spectra: those modeling single cells explicitly often rely on vertex-based approaches and are limited to dozens of cells (Alt, Ganguly, and Salbreux 2017; Misra et al. 2016; Benoît Aigouy et al. 2010; Le Garrec, Lopez, and Kerszberg 2006). To capture the large features spanning thousands of cells, one typically turns to elastic models where AB polarity is implicit and epithelia is presented as a 2D elastic sheet (Hannezo, Prost, and Joanny 2014; Etournay et al. 2015; Hufnagel et al. 2007; Nagai and Honda 2009; Aliee et al. 2012; Nagai and Honda 2001).

We developed a theoretical approach that, with only a few parameters, bridges cellular and organ scales by integrating both types of polarity. A main difference to earlier approaches is that a cell’s movement is coupled to how its AB polarity and PCP are oriented relative to each other and relative to neighbor cell polarities. In other words, in our approach the adhesion strength between neighbor cells is modulated by the orientation of their polarities. We find that polarity enables complex shapes robust to noise but sensitive to changes in initial and boundary constraints, thus supporting that morphological stability and diversity are emergent properties of polarized cell populations. Lumens, folds, and stable tubes emerge as a result of energy minimization. We make testable predictions on morphological transitions in pancreatic organoids, tubulogenesis, and sea urchin gastrulation. Our approach illustrates the evolutionary flexibility in the regulatory proteins and networks, and suggests that despite differences in proteins between organisms, the same core principles may apply.

## MODEL

There are three key elements that allow us to bridge the scale from cellular level to macroscopic stable morphologies.

### (1) Cells are approximated by point particles

Cellcell adhesion is modeled by repulsive and attractive forces acting between cell centers. This allows a substantially gain in computation time compared to vertex based models where cell surface adhesion are explicitly considered (Alt, Ganguly, and Salbreux 2017). The potential for pairwise interaction between two interacting neighbors, *i* and *j*, separated by distance *r*_*ij*_ is

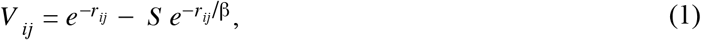

where the first term corresponds to repulsion, and the second term to attraction (see Figure 2A). For a pair of non-polar cells the strength of attraction *S* = 1. *β* > 1 is the parameter that sets how much longer the attraction range is compared to repulsion. We set *β* = 5 throughout the paper, but our results and conclusions are consistent for smaller *β*. The main results are also not sensitive to the exact choice of the potential, thus for example the higher power in the exponential,

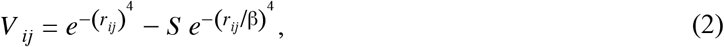

give qualitatively similar results (see Figure S2A). The potential energy of a cell is the sum of pairwise neighbor interactions

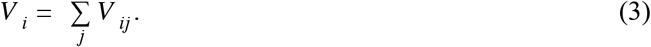

**Figure 2.**
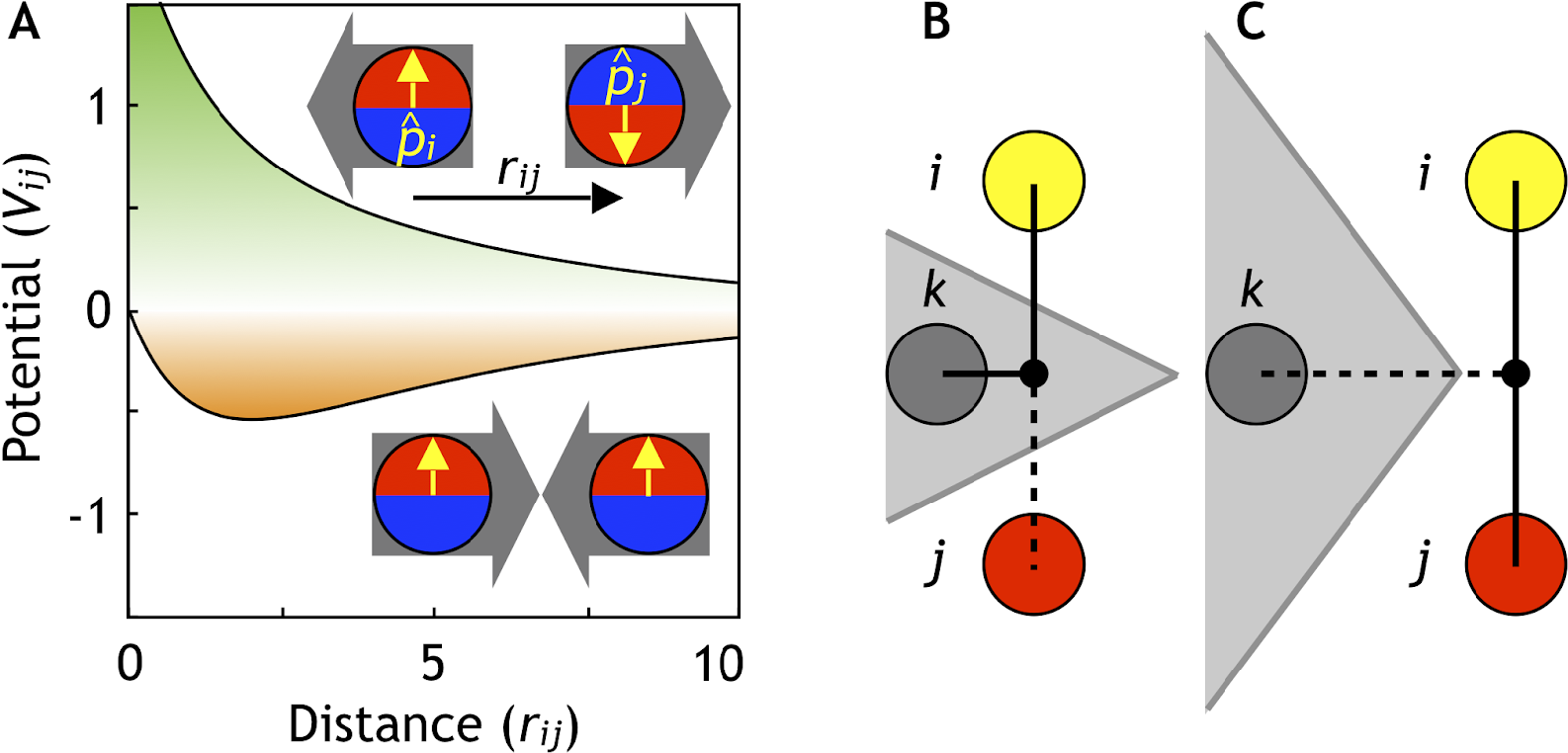
Cells are modeled as interacting particles with a polarity-dependent potential. **(A)** Potential between two interacting cells with apical-basal polarity (see Equation 6). Cells repulse when polarities are antiparallel (top/green part) and attract when they are parallel (orange/ bottom part). **(B-C)** Two cells interact only if no other cells block the line of sight between them. **(B)** Cell *i* and *j* do not interact if *ij*’s midpoint (black dot) is inside of the Voronoi diagram for cell *k* (shaded in grey). **(C)** Cell *i* and *j* interact because cell *k* is further away than the distance *ij*/2 and *ij*’s midpoint therefore lie outside of cell *k*’s Voronoi diagram. In the related Figure S2A-D, we test the sensitivity of our model to the details of the potential and neighborhood assignments. In Figure S3, we relate changes in cell shapes to the model components, and in Figure S4 (Movie 1), we illustrate how altering polarity affects the dynamics of the systems with two and six cells.

### (2) Cells interact with (a subset of Voronoi) neighbors

Interacting neighbors of cell *i* are selected from a subset of cells sharing a Voronoi surface. The subset is limited to the nearest neighbors *j* which are closest to the midpoint between *i* and *j* (Figure 2B-C). This constrain effectively corrects for the finite volume associated with point particles and assures that two cells will not interact if the line of sight between their centers is separated by a surface of a third cell. Without this constraint, the macroscopic morphologies collapse. However, our results are robust to replacing the line of sight constraint with full Voronoi and a cut-off distance for attraction force (Figure S2B).

### (3) Cell-cell adhesion depends on the orientation of polarity

To capture directional adhesion, we set the strength of attraction, *S*. to be dependent on the relative orientation of the polarities in each of the cells. We assume that AB polarity and PCP are orthogonal, and that the polarities of one cell align with the polarities of it’s neighbor cells. Mathematically, we introduce unit vectors, 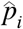, and 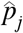, representing AB polarity, and 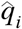 and 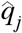 representing PCP for cell *i* and *j*, respectively. We set

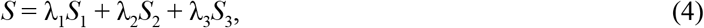

where *λ*_1_, *λ*_2_ and *λ*_3_ are the strengths of the different polarity terms. We require that

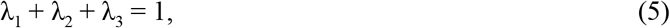

to satisfy the constraint that perfectly aligned cells always have a steady state distance of 2 cell radii (for *β* = 5). To capture that in an epithelial sheet, AB polarities align parallel to each other and tight adherens junctions form in the plane perpendicular to AB polarity, we introduce the quadruple product

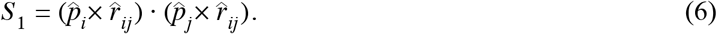

This makes two interacting cells with AB polarity maximally attracted (*S*_1_, = 1) if the two apical sides are next to each other. On the other hand, if apical side of one cell is next to the basal side of another cell, the two cells will be maximally repulsing (*S*_1_ = −1), see Figure 2A.

In case of planar polarization, we define

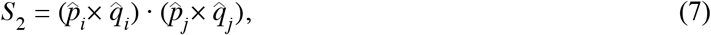

which makes the attraction maximal if the PCP of two cells are parallel to each other and perpendicular to their AB polarities. In addition, we assume that similarly to AB polarity, two cells with PCP are maximally attracted if their PCP are parallel and cells have the same kind of pole (e.g. Vangl-enriched) next to each other,

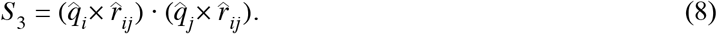

We later show that this assumption makes neighbor exchange on a sheet possible and results in CE. However, unlike with tight junctions, preferred directional adhesion with PCP is not as well established. Cells adhere to each other by membrane proteins assembled in adherens junctions just below the apical surface. Both proteins regulating adherens junctions, e.g. Smash (Beati et al. 2018), as well as adherence proteins forming adherens junctions can be planar polarized, e.g. Bazooka, E-cadherins (Simões et al. 2010; Tamada, Farrell, and Zallen 2017; Levayer and Lecuit 2013; Warrington, Strutt, and Strutt 2013; Benoit Aigouy and Le Bivic 2016). These indirectly support our assumption of anisotropic, planar polarized adhesion.

The motion of the cells and their polarities are calculated assuming overdamped dynamics

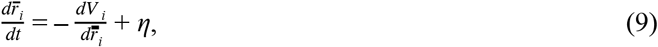

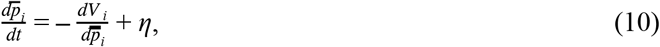

and

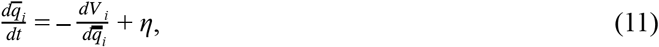

where the 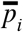 and *q̅*_*i*_ differentiation takes into account the rotation of polarity vectors, and *η* is a random uncorrelated Gaussian noise. In practice, we implemented the model in a MatLab script (available in the supplementary material), where we use the Euler method. We perform the differentiation along the polarity by differentiating along all three cartesian coordinates (see Model in the supplementary material). After each time step, we normalize the updated polarity vectors. The above differentiation does not include the change in partners when neighborhood changes. This is treated as a non-equilibrium step where potential energy can increase (Equation 3). Biologically, this is similar to cells spending biochemical energy as they rearrange their neighborhood.

The point particle approximation has been utilized earlier for modeling non-polar cell adhesion in early blastocyst (Krupinski, Chickarmane, and Peterson 2011), slug formation in amebae (Dallon and Othmer 2004), and PCP organization in primitive streak formation (Newman 2008). The main novelty of our approach is the dynamical coupling of cell positions and polarity orientations (Equation 6–8).

#### Model implications

One of the implications of the coupling between position and polarity is that in a sheet of cells, turning AB polarity in one cell will cause a force on its neighbors. In case of two cells (Figure S4 and Movie 1) the pair relaxes the imposed stress by rotating both the polarity and their positions. In biological terms, turning AB polarity in one cell (e.g. by apical constriction, illustrated in Figure S3) of an epithelial sheet will induce bending of the sheet as is the case with bottle cells in invagination.

The present formulation of PCP has several implications. First, we restrict the effects of PCP to directed (anisotropic) cell-cell adhesion and do not consider its other possible roles, in e.g. asymmetric cell differentiation, thus primarily focusing on its role on CE. Second, in our current formulation AB polarity and PCP influence each others orientation on equal footing (Equation 7). PCP, however, is typically constrained to the apical plane and thus is expected not to influence the orientation of AB polarity. Disabling PCP’s effect on AB polarity (see Methods) does not influence our main results on tube formation and gastrulation. However, the symmetry in polarities is appealing for its simplicity and is indirectly supported by the following experimental observations: First, cells can acquire PCP without AB polarity present (Baer, Chanut-Delalande, and Affolter 2009; Zorn and Wells 2009). Second, proteins required for AB polarity can be planar-polarized (Warrington, Strutt, and Strutt 2013; Benoit Aigouy and Le Bivic 2016; Beati et al. 2018; Choi and Sokol 2009; Dollar et al. 2005; Kaplan and Tolwinski 2010). Third, changes in cell shapes during invagination (e.g. sliding of adherens junctions and formation of bottle cells) are regulated by PCP in neural tube closure (Ossipova et al. 2014; Nishimura, Honda, and Takeichi 2012; Kinoshita et al. 2008), gastrulation in *C. elegans* (Lee et al. 2006), sea urchin (Croce et al. 2006), and *Xenopus* (Choi and Sokol 2009). These changes in cell shape effectively reorient AB polarity (Figure S3).

## RESULTS

We have recently introduced effective representation of AB polarity, and showed that it is sufficient for capturing spherical trophectoderm in the early blastocyst (Nissen et al. 2017). Expanding on that work, we here explore how AB polarity supports diverse yet stable and complex morphologies.

### Stable complex shapes emerge from randomly polarized cell aggregates

Adult organismal shapes are stable over long time, maintaining sizes and relative positions of lumens and folds, despite continual local damages and cell renewal. To test if cellular polarity could enable such stability in time and to random local perturbations, we first performed a series of tests with AB polarized cells (Figure 3 and Movie 2).

**Figure 3.**
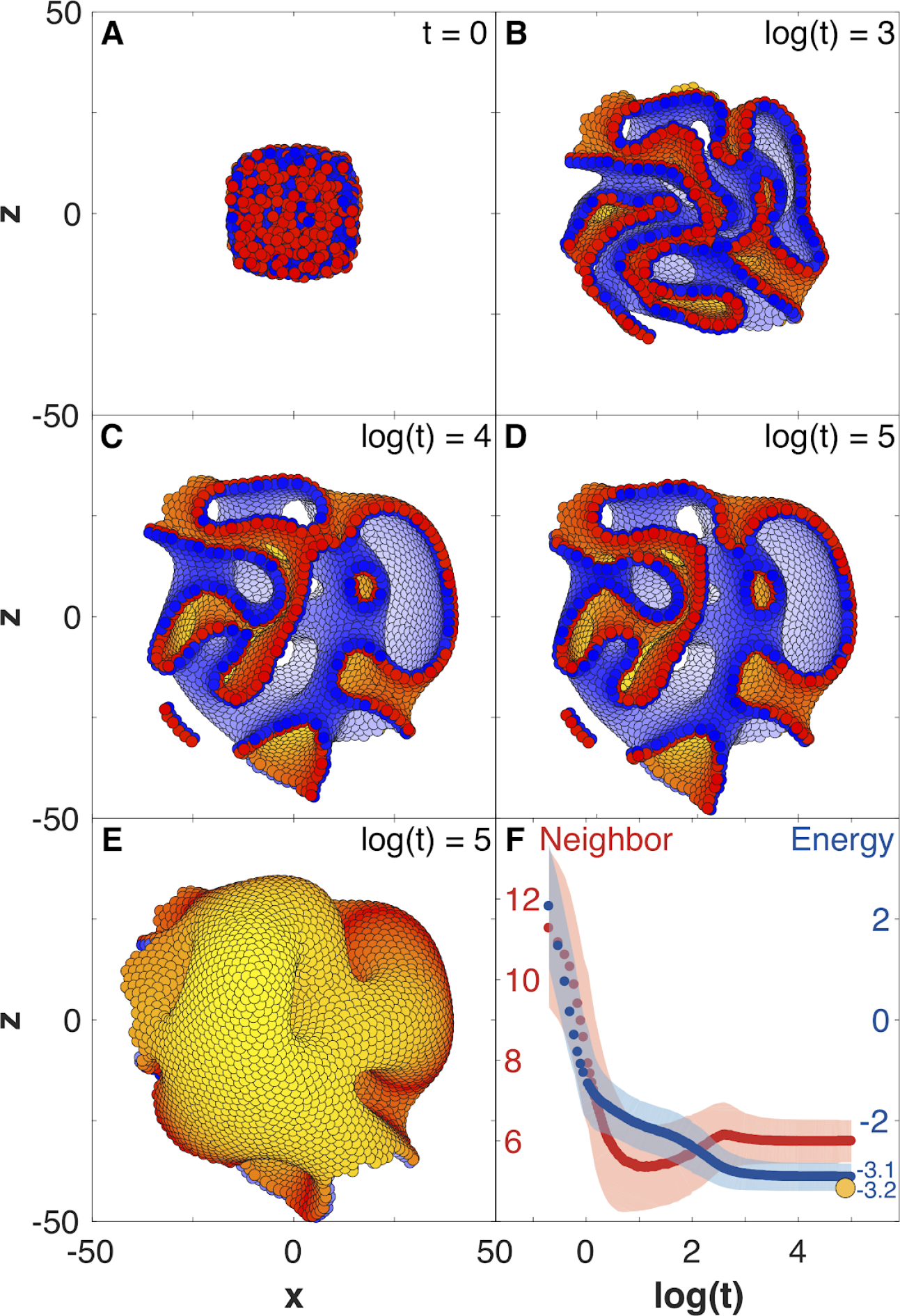
Development of 8000 cells from a compact aggregate starting at time 0. **(A)** Cells are assigned random apical-basal polarity directions and attract each other through polar interactions (see Equation 6). **(A-D)** Cross section of the system at different time points with red and blue marking two opposite sides of the polar cells. Cells closest to the viewer are marked red/blue, whereas cells furthest away are yellow/white. **(E)** Full system at the time point shown in (D). **(F)** Development of the number of neighbors per cell (red) and the energy per cell (blue), as defined by the potential between neighbor cells in Figure 2. Dark colors show the mean over all cells while light-shaded regions show the cell-cell variations. The yellow dot marks the energy for a hollow sphere with the same number of cells. See Movie 2 for full time series. In Figure S2E-G and Figure S5, we study how the final morphology depends on noise. In Figure S6, we show how the outer surface self-seals, and that the shape is maintained when cells divide.

When starting a bulk of cells with AB polarities pointing randomly, an initial rapid expansion (Figure 3A-C) stabilizes into a complex morphology of interconnected channels (Figure 3C-E). The shape remains unchanged for at least 10 times longer than the initial expansion (Figure 3C-E). The stability of the shape is illustrated by the time evolution of the average energy per cell (Figure 3F) that after an initial fast drop converges to a constant value. As expected, this value is higher than the energy of a hollow sphere (yellow dot in Figure 3F) - a configuration obtained if we start with radially, instead of random, polarized cells and preserve radial polarization at all times. The observed behaviour is not sensitive to the shape of the potential (Figure S2A) but is sensitive to how the neighborhood is defined (Figure S2B-D). Rerunning the simulation in Figure 3 with different initial conditions results in a different stable shape (Figure S2E and Figure S5).

The macroscopic features of the shapes are robust to noise (Figure S2E-F and Figure S5). While the shapes emerging under high and low noise are not identical, the relative position and sizes of the majority of channels and lumens are preserved. The changes caused by noise stem from perturbations during initial expansion stage. If the same level of noise is applied after the system reached stable state, after time *t* = 10000, noise does not cause any major macroscopic changes (Figure S2G). The obtained shapes have self-sealing features, as an initial cut and unwrapping of a section of a surface refolds and seals back into the original morphology (Figure S6A-C). Furthermore, the shapes (Figure 3D-E) are also robust to overall growth (Figure S6D-F) retaining the same macroscopic features, just scaled to a larger size. Robustness to noise and cell proliferation further support the link between polarity and stability of morphologies, e.g. organ shapes, as they expand from infant to adult.

### The final shapes are robust to noise but sensitive to initial and boundary conditions

The orientation of polarities in a subpopulation of cells may be set by the environment that the cells are embedded in, e.g. signaling molecules deposited into extracellular matrix can influence orientation of the AB polarity (Overeem, Bryant, and van IJzendoorn 2015) or signals from neighboring cells of a different type can orient PCP (Chu and Sokol 2016). We will refer to these constraints as boundary conditions.

To investigate sensitivity to boundary conditions, we consider three cases where polarities are fixed at all times and point either radially out from the center of mass (Figure 4A), radially out from a central axis (Figure 4B), or pointing away from a central plane (Figure 4C). As anticipated, the difference in symmetries of boundary conditions results in a sphere, a cylinder, or two parallel planes (Movie 3). At the same time, in these symmetric cases the differences in initial conditions but without imposed boundary conditions are not sufficient to generate different structures; they all converge to the nested “Russian doll”-like hollow spheres (Figure 4D). In development, this highlights the importance of the neighboring tissues for defining boundary conditions.

**Figure 4.**
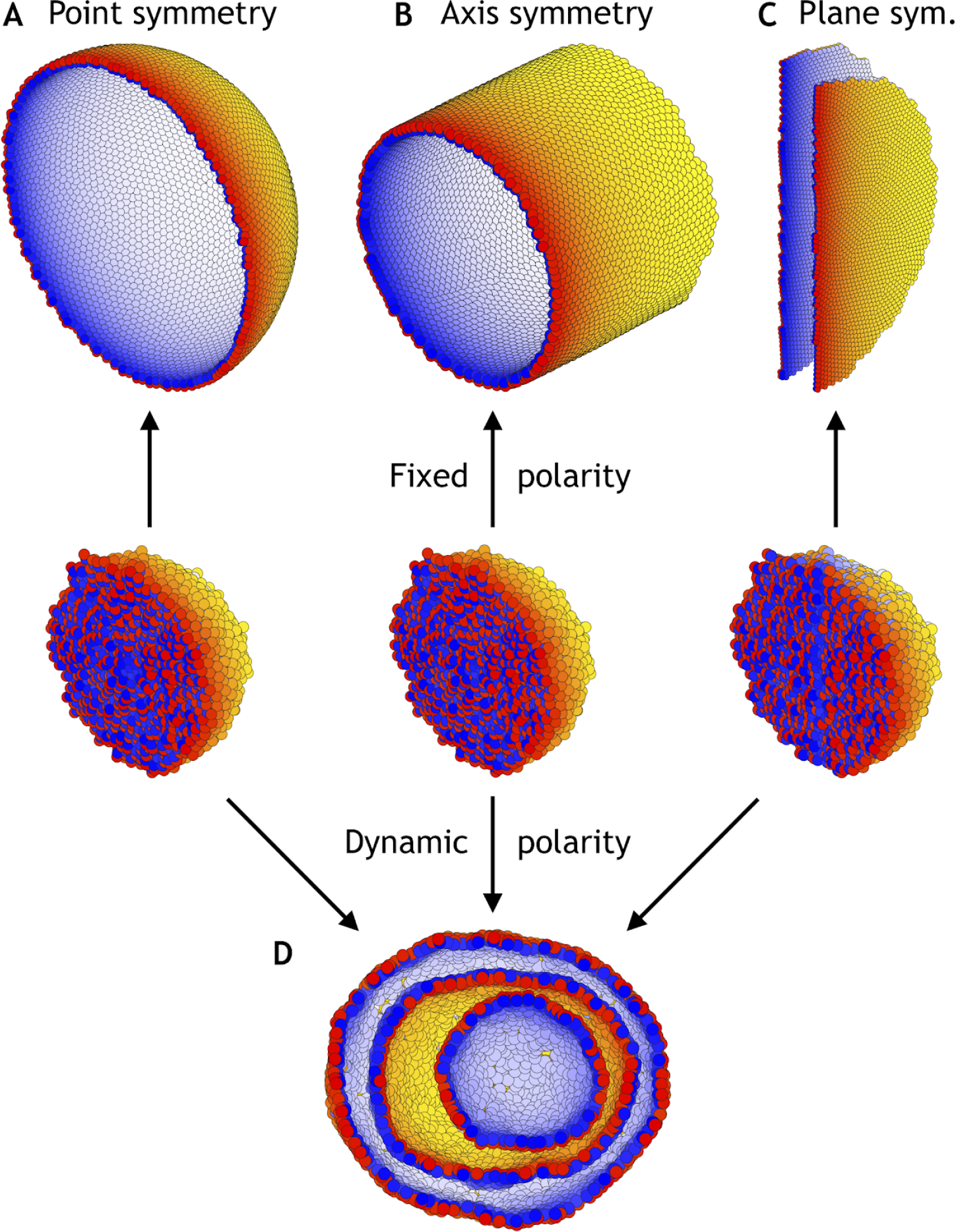
Different morphologies can be obtained by varying boundary conditions (Movie 3). **(A)** A hollow sphere emerges if polarities are fixed and initially point radially out from the center of mass. **(B)** A hollow tube is obtained if polarities point radially out from a central axis. **(C)** Two flat planes pointing in opposite directions are obtained if polarities point away from a central plane. **(D)** For all three initial conditions (A-C), if the polarities are allowed to change dynamically and the noise is high (*η* = 10° compared to *η* = 10^−1^ in A-C), the resulting shape consists of three nested “Russian doll”-like hollow spheres that will never merge due to opposing polarities. In contrast to the random initial condition in Figure 3, the initial conditions in (D) are symmetric.

Our results thus support the idea that polar adhesion enables stable and robust macroscopic shapes. The closest biological parallels would be the complex luminal morphologies emerging in reaggregation experiments on e.g. *Hydra* (Seybold, Salvenmoser, and Hobmayer 2016) or *in vitro* culture of purjunkie brain cells (Muguruma et al. 2015). Together with our simulations, these experiments highlight how stable and complex morphologies can develop in non-proliferating populations from cell rearrangements alone.

### Folding by pressure or rapid proliferation result in different fold-m orph ologies

Transitions from spheres to folded shapes are ubiquitous in development. Folds are an important part of *in vivo* organ development, and the composition of cell types in the folded organoids is closer to that in real organs (Greggio et al. 2013). To date, it is unclear what drives the transition from spheres to folded lumens. One possibility is that it is driven by the mechanical properties of the matrigel that effectively may place the growing organoid under pressure. Alternatively, data from 3D brain organoids suggests that the rapid cell proliferation leads to the emergence of surface folding (Y. Li et al. 2017).

The simplicity of our tool allows to explore both of these scenarios. To model dividing cells, we pick a random cell from the entire population and introduce a new daughter cell with inherited polarity direction placed in a random location a half cell radius away from the mother cell. This event introduces dynamic perturbation by locally increasing cell density and requires some time to relax back to equilibrium. If proliferation is slow, and the time between two cell divisions anywhere in the system is longer than relaxation time of the whole system (the time it takes to reach equilibrium), the system approaches global equilibrium and will expand as a sphere. However, if proliferation is increased, the system will be pushed out of equilibrium and folds will emerge (Figure 5A, Methods). In more quantitative terms, our forces are such that a single cell can move up to 0.2 cell diameter per time unit. For cells that divide every 1000 time units, the transition to non-equilibrium buckling happens when the system has grown to about 5000 cells (Movie 4).

**Figure 5.**
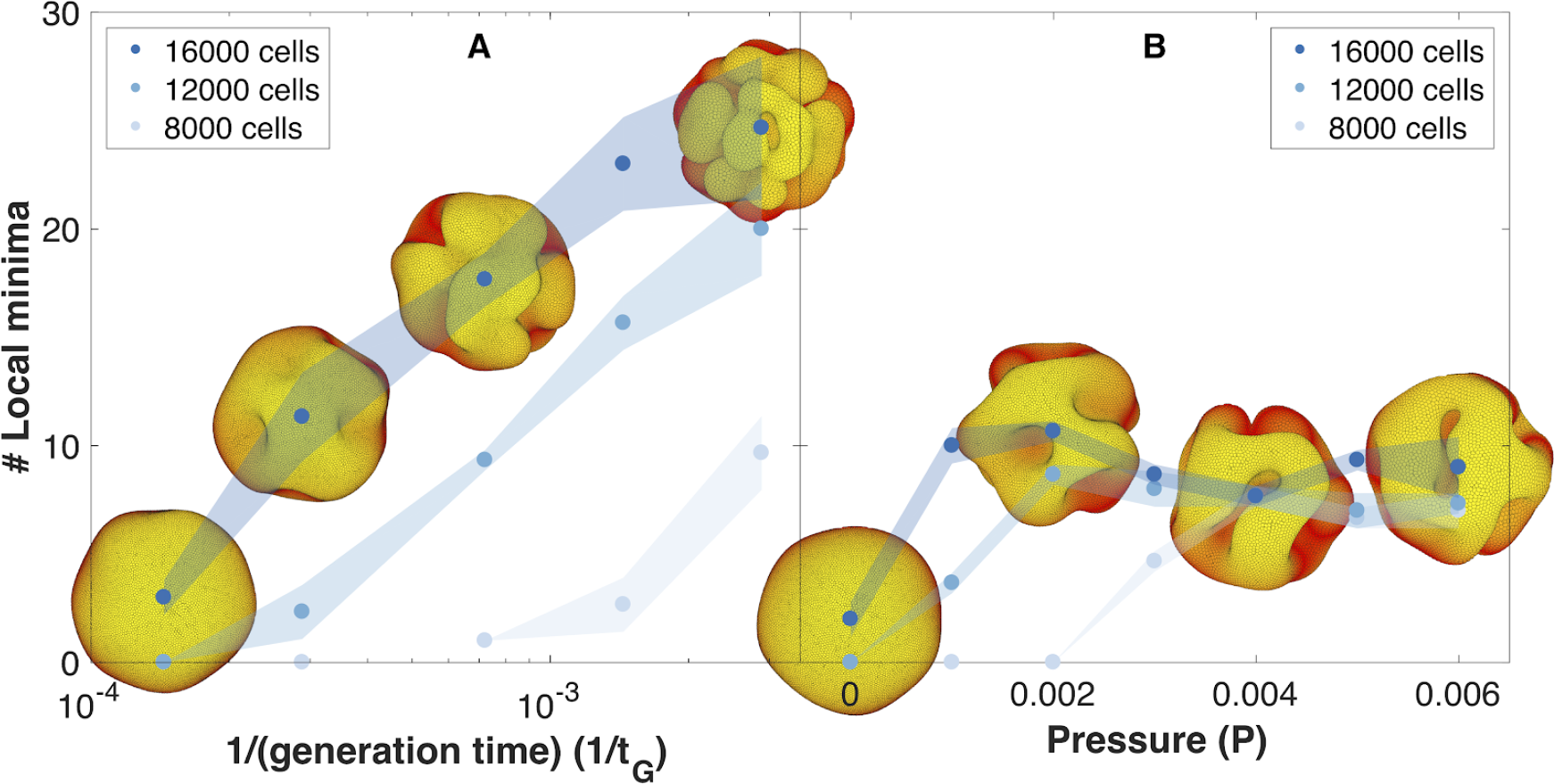
The number of complex folds in a growing organoid depends on the generation time and the pressure from the surrounding medium (Movie 4). **(A)** Number of local minima as a function of 1/(generation time), *t*_*G*_^−1^. *In silico* organoids grow from 200 cells up to 8000, 12000, or 16000 cells with different generation times and no outer pressure. **(B)** Number of local minima as a function of pressure, *P. In silico* organoids grow to the same size with the same 1/(generation time), *t*_*G*_^−1^ = 1.4·10^−4^ but different outer pressure. The images illustrate the 16.000 cells stage. Blue dots mark the average, while light shaded regions show the SEM based on triplicates. See also Figure S7 for additional measurements on the differences between rapid growth and pressure.

As cells divide faster, our simulations predict a transition from a smooth spherical shell to an increasingly folded structure with multiple pronounced folds, in line with the observation of brain organoids proliferating at different rates (Y. Li et al. 2017). In comparison with the model for cortical convolutions by (Tallinen et al. 2016) in which folding is a result of expanding cortical sheet adhered to the non-expanding white matter core, our mechanism does not require a bulk core. Instead folds emerge in a fast expanding sheet when the growth is faster than the global relaxation to dynamical equilibrium.

While we find that the external pressure is not necessary for folding, pressure alone can also drive folding (Figure 5B, Methods). However, this scenario contradicts the observation that pancreatic organoids can grow as spheres or folded morphologies in gels with the same stiffness but different media composition (Greggio et al. 2013).

In principle, both scenarios may contribute to folding, but visually the fold morphologies are different. To differentiate between the two, we have quantified the final folded structures in terms of their local minima (Figure 5, Methods, see also Movie 4). Our simulations predict that in the pressure driven case, the number of local minima will reach an upper limit as organoids increase in size (Figure 5B). In the case of out-of-equilibrium proliferation, new folds can continue forming as organoids grow (Figure 5A). Increased proliferation causes more and shallower folds. These folds are different than obtained with pressure which causes a shape with fewer but deeper minima. Quantitatively, both the depth and the horizontal extension of the folds are about double as large with pressure than with growth-induced folding (Figure S7).

### PCP enables convergent extension and robust tubulogenesis

Despite the numerous evidence supporting the role of PCP in tubulogenesis, it remains unresolved whether oriented cell division or the extent of CE controls tube length and width (Karner et al. 2009; Carroll and Yu 2012). It is also debated if it is important for the tubes to maintain regular shape, or if it is only important for tube initiation and growth (Kunimoto et al. 2017).

The simplicity of our approach allows us to address these questions by introducing cell-cell interactions through PCP. This term favours front-rear cell alignment in the interaction potential with only two additional parameters: the strength of the orientational constraint of AB polarity with respect to PCP, *λ*_2_, and the strength of PCP, *λ*_3_, (see Equations 4–8). For simplicity, we focus on the stability (ability to maintain regular diameter over time) and tube morphogenesis in systems without cell division.

Inducing PCP in a spherical lumen leads to two significant events. First, independent of initial orientation, after some transient time PCP becomes globally ordered and point in direction parallel to an emerging equator that self-organizes around the sphere (inset in Figure 6). This arrangement has the lowest energy. Second, cells start intercalating along the axis perpendicular to PCP orientation, gradually elongating the lumen (Movie 5). During intercalations, cells exchange their neighbors through Tl-like transitions as reported experimentally, Figure S10 (Nishimura, Honda, and Takeichi 2012; Sanchez-Corrales, Blanchard, and Röper 2018). The intercalations along the axis continue until the force balance between AB polarity and PCP is restored at a new equilibrium. Thus, our model predicts that the strength of PCP (*λ*_3_) relative to AB polarity (*λ*_1_) determines the width and the length of the tube (Figure 6). We obtain similar results if we constrain PCP to always remain in the apical plane and thus does not allow PCP to reorient AB polarity (Figure S8, Methods). Note, that this result is very different in nature from the tube presented in Figure 4B as both AB polarity and PCP can now reorient in each cell at any time.

**Figure 6.**
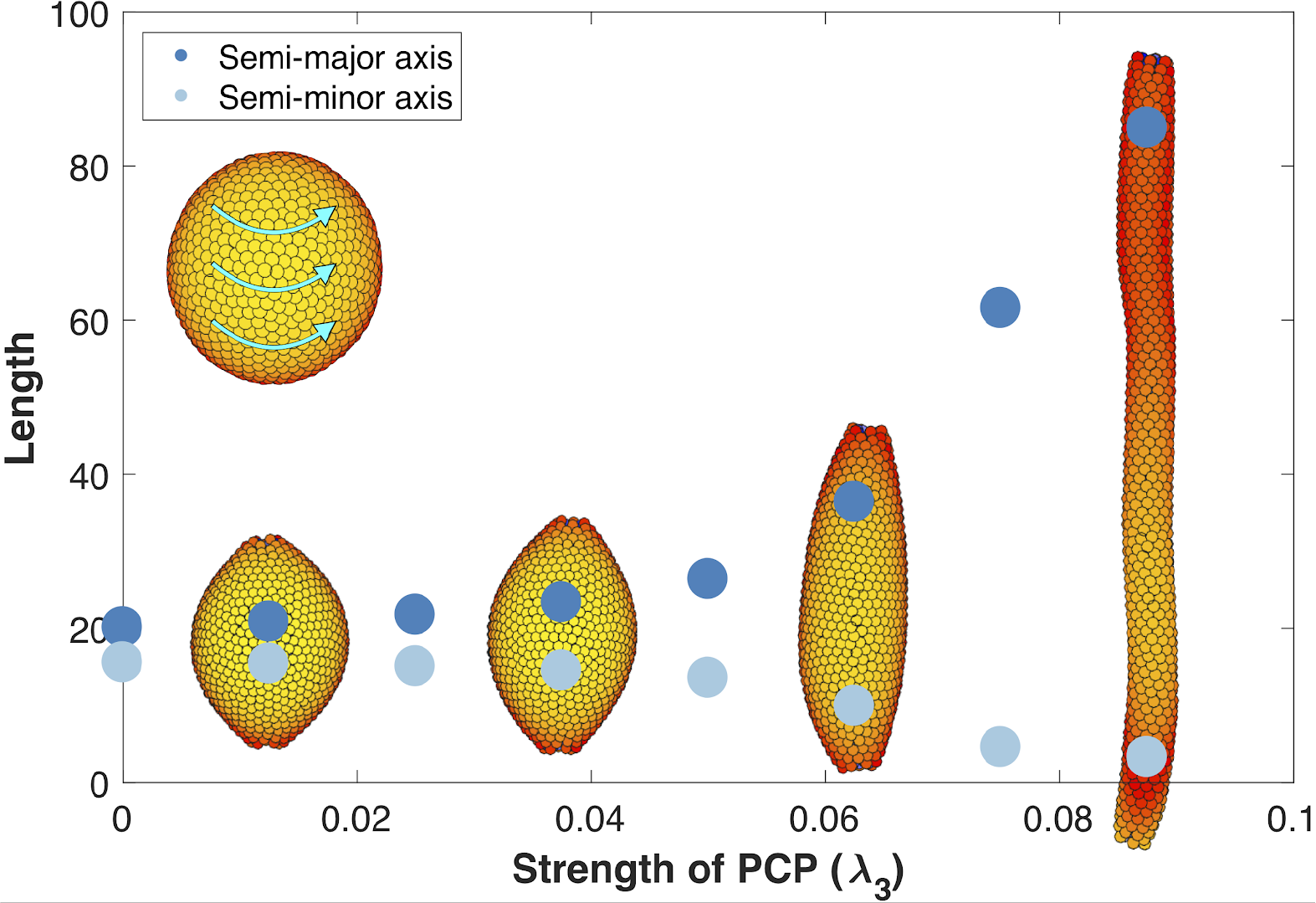
The length and width of tubes are set by the strength of planar cell polarity (PCP, *λ*_3_). For each value of *λ*_3_, we initialize 1000 cells on a hollow sphere with PCP whirling around an internal axis (PCP orientation marked by cyan arrows in the top-left inset). Semi-major axis (dark blue) and semi-minor axis (light blue) are measured at the final stage (Methods). Images show the final state. Throughout the figure, *λ*_2_ = 0.5 and *λ*_1_ = 1 − *λ*_2_ − *λ*_3_. The animated evolution from sphere to tube is shown in Movie 5. See also Figure S8 where we show that tubes also form when we disable the direct influence of PCP on apical-basal polarity, and Figure S9 where we vary the degree of PCP along the axis of the tube. In Figure S10, we show that cell intercalations result in experimentally reported T1 neighbor exchanges during convergent extension.

These results support the observations that stable tubes can emerge without cell proliferation. In addition, when first the tube is formed, loss of PCP does not lead to cyst formation as recently shown by Kunimoto et al. (2017). However, localized cysts could result if the lumen is initialized with varying strength of PCP along the axis (Figure S9).

### Two polarities are sufficient to explain major features of sea urchin gastrulation

Currently invagination in neurulation and gastrulation is understood and quantitatively modeled as a process driven by changes in cell shapes or the mechanical properties of cells with AB polarity (Rauzi et al. 2013; Tamulonis et al. 2011; Misra et al. 2016; Hočevar Brezavšček et al. 2012). This process is often assumed to be driven by apical constriction and decoupled from the eventual tube formation and elongation. However, emerging data suggests that PCP drives both invagination and tube elongation (Nishimura, Honda, and Takeichi 2012; Croce et al. 2006; Long et al. 2015) likely because apical constriction is controlled by PCP (Ossipova et al. 2014; Nishimura, Honda, and Takeichi 2012; Croce et al. 2006) and cell intercalations, similar to those in CE, contribute to invagination (Nishimura, Honda, and Takeichi 2012; Rembold et al. 2006; Sanchez-Corrales, Blanchard, and Röper 2018).

To probe the limits of our approach, we investigated if AB polarity, PCP, and boundary conditions reminiscent of posterior organizer (Loh, van Amerongen, and Nusse 2016) are sufficient to recapitulate the main stages of sea urchin gastrulation: invagination, tube formation, elongation (by CE), and finally merging of the gastrula tube with the pole opposite to invagination site.

The current understanding is that sea urchin gastrulation consists of primary invagination, driven by swelling of the inner layer of extracellular matrix beneath invaginating cells (Lane et al. 1993), and formation of a ring of bottle cells due to apical constriction (Kimberly and Hardin 1998), and secondary invagination where tube elongates due to CE (Lyons, Kaltenbach, and McClay 2011). PCP is necessary for both invagination, possibly through its effect on apical constriction in bottle cells (Nishimura, Honda, and Takeichi 2012), and tube extension (Croce et al. 2006).

Motivated by these observations, we set boundary condition such that PCP of the invaginating cells are oriented around the anterior-posterior (top-bottom) axis, and are always in the apical plane. This constrain on PCP orientation allows for CE. While this particular configuration is not documented, it is consistent with observed effects of WNT orienting PCP within the apical plane (Humphries and Mlodzik 2017). Second, we simulate the combined effect of bottle cells (Figure S3) and bending by swelling of the extracellular matrix by applying an external force, *F*, on AB polarity (see Methods). This force gradually reorients AB polarity away from the anterior-posterior axis, thus leading to bending of the epithelial sheet. The effect is maximal for cells closest to the anterior-posterior axis. This external force is a phenomenological description aiming at capturing the observed effects of how change in AB polarity results in tissue bending and does not aim at capturing mechanisms driving the reorientation of AB polarity.

As a result cells start to rearrange, the bottom flattens (Figure 7B) and bends inward (Figure 7C). Subsequently, the CE-driven by PCP causes the invaginated cells to rearrange, tube elongates, and merges with the top of the sphere (Figure 7F and Movie 6). In line with experimental observations, the tube elongates due to cells moving into the tube (Martik and McClay 2017).

**Figure 7.**
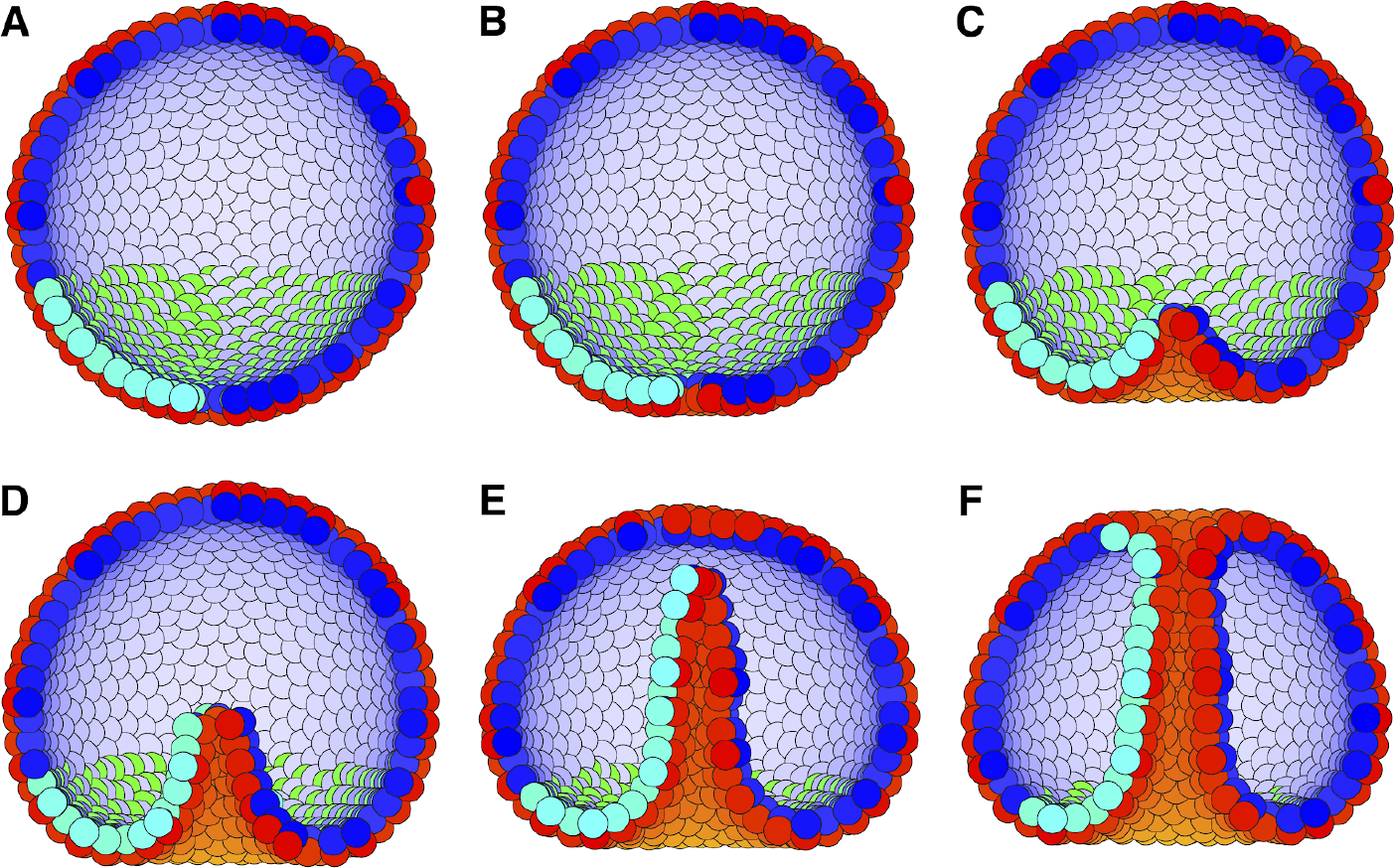
External constraints on apical-basal (AB) polarity and planar cell polarity (PCP) can initiate invagination and drive gastrulation in sea urchin. **(A)** The lower third of the cells in a blastula with AB polarity (apical is blue-white, basal is red-orange) pointing radially out acquire PCP (cyan-green) in apical plane pointing around the anterior-posterior (top-bottom) axis (as in the inset to Figure 6). **(B)** Flattening of the blastula and **(C)** invagination occur due to external force reorienting AB polarity (Methods). **(D-E)** Tube elongation is due to PCP-driven convergent extension and **(F)** merging with the top of the blastula happens when the tube approaches the top. Throughout the simulation *λ*_1_ = 0.5, *λ*_2_ = 0.4, and *λ*_3_ = 0.1 for the lower cells while the top cells have *λ*_1_ = 1 and *λ*_2_ = *λ*_3_ = 0. For full time dynamics see Movie 6. In Figure S11 we consider alternative scenarios of sea urchin gastrulation and and neurulation.

The contribution of extracellular matrix swelling is specific to sea urchin invagination. As bottle cells alone are sufficient to drive invagination in other systems, we have tested a cell-intrinsic scenario where AB polarities of bottle cell neighbors prefer to be tilted towards each other (by e.g. modifying the potential in Equation 6 to capture AB polarities as in Figure S3, unpublished results). This also resulted in successful invagination.

Several observations suggest that radial intercalation movements towards the center of invagination may drive tissue bending (Sanchez-Corrales, Blanchard, and Röper 2018; Panousopoulou and Green 2016; Rembold et al. 2006; Chung, Kim, and Andrew 2017). When tested in our model, PCP alone (omitting external or cell-intrinsic reorientation of AB polarity) results in exvagination and tube elongating outside of the sphere (this is more energetically favourable than extending inwards). A similar, exogastrulating, phenotype was observed in a PCP mutant (Long et al. 2015). While the similarity may be accidental, it is possible that this mutation abrogates PCP-driven apical constriction (Ossipova et al. 2014; Nishimura, Honda, and Takeichi 2012) and since cellular apices face outside, abrogating apical constriction eliminates the bias in direction of tube formation. We have also tested the consequences of apical constriction (reorienting AB polarity) in the model without PCP and while this was sufficient for invagination, the cavity remained spherically symmetric and failed to form a tube.

## DISCUSSION

Despite the stunning diversity and complexity of morphologies, the same concepts seem to emerge across organismal development. One of them is the link between local, cellular, and global, organs/whole organism, symmetry breaking. We know, from experimental and, to a lesser degree, theoretical work that cellular polarity is essential for forming axis, complex folded sheets, and interconnected tubes (Figure S1). What we do not know is why are these shapes so stable, and where do the differences between species and organs come from. To understand the differences, we typically compare genes or gene regulatory networks, thus limiting our understanding to analog processes within a few related species.

### Phenomenological description bridges cell polarity to macroscopic morphologies

To address the origins of morphological diversity and stability across species and organs, we focused on a phenomenological description of polarized cell-cell interactions. This allowed us to bridge local single cell symmetry breaking events to global changes in morphologies spanning tens of thousands of interacting cells. With this tool at hand, we find that with only a few parameters, we can recapitulate the two global symmetry breaking events: formation of epithelial sheets and folds by cells with AB polarity, and emergence of global axial symmetry (tubes) among cells with PCP.

Remarkably, our results show that interactions among AB polarized cells lead to stable morphologies, that after initial relaxation remain indefinitely in their final configuration. The morphologies are robust to noise, growth, and local damage (Figure S6). These results may explain how organs and embryos preserve their architecture while growing. Polar cell-cell interactions not only provide clue to the morphological stability, but also point to a simple explanation to the origin of the diversity. We find that the exact morphological details are defined by initial conditions, e.g. initial positions and orientation of polarities, and boundary conditions, e.g. polarities restricted to certain direction for a fraction of the cells. It is thus tempting to speculate that diverse shapes do not require multiple interacting morphogen gradients, but simply can be a result of differences in initial and/or boundary conditions: as for example presence of yolk cells at start and boundary constraints by vitelline membrane (Schierenberg and Junkersdorf 1992; T. Wu et al. 2010).

The diversity of shapes and forms is further enriched by a second symmetry breaking event, PCP, oriented perpendicular to AB polarity. Within our phenomenological framework, addition of PCP component is simple, and requires only two additional parameters: one favouring perpendicular orientation of AB polarity and PCP within a cell, and another, favouring parallel PCP alignments between neighbor cells. These constraints are the coarse-grained representation of the well-established experimental and computational results on intracellular symmetry breaking events and global ordering of planar polarities mediated by cell-cell coupling (Le Garrec, Lopez, and Kerszberg 2006; Amonlirdviman et al. 2005; Wang, Badea, and Nathans 2006). The first constraint allowed formation of axial symmetry and in combination with AB polarity, stable tubes, with length and diameter remaining constant with time. The second constraint resulted in cell rearrangements and intercalations consistent with the cell-autonomous CE typically associated with PCP. The patterns of neighbor exchanges during cell-rearrangements are in line with the ubiquitous T1 exchanges through formation and resolution of four cell vertex, Figure S10 (Nishimura, Honda, and Takeichi 2012; Sanchez-Corrales, Blanchard, and Röper 2018). The mechanism of the CE in our model is in line with the results by “filopodia tension model” where elongated structures of many cells emerge from local cell-cell interactions in a direction defined by PCP (Belmonte, Swat, and Glazier 2016). The presented formulations of our model captures only some of the known events contributing to CE and does not include PCP-driven changes in cell shape mediated by e.g. apical constriction (Nishimura, Honda, and Takeichi 2012) or contribution of external forces (Lye and Sanson 2011).

Combining AB polarization and a local induction of PCP in a subpopulation of cells was sufficient to obtain main stages of sea urchin gastrulation: invagination, tube formation, and elongation through CE as well as merging of the tube with the animal pole at the top of the blastula.

The existing *in silico* models treat invagination and CE-driven tube elongation as independent processes (Figure S1). Recent data, however, suggests that multiple mechanisms (intracellular apical constriction, intercellular directed cell division and cell intercalations, and supracellular actomyosin cables) act simultaneously and contribute to both invagination and tubulogenesis (Chung, Kim, and Andrew 2017; Nishimura, Honda, and Takeichi 2012; Ossipova et al. 2014). Within our approach apical constriction (modeled as reorientation of AB polarity) and CE can act in parallel (Figure 7). It will be interesting to parallel recent experimental work (Chung, Kim, and Andrew 2017) and computationally investigate how a combination of intra-, inter- and supracellular mechanisms contribute to the robustness of tubulogenesis, and to what extent the model can capture the range of observed phenotypes.

It has been proposed that the above mechanisms may all be coordinated by PCP (Nishimura, Honda, and Takeichi 2012). Besides the reported molecular links, a simple logic suggests that these mechanisms can not be isotropic as in this case the initial bending will result in spherical structures. Thus, apical constriction, cell intercalations, and actomyosin cables have to be anisotropic (planar polarized) in directions consistent with the eventual tube orientation. This anisotropy is reported for both “wrapping” tubes forming parallel to the epithelial plane, e.g. neurulation (Nishimura, Honda, and Takeichi 2012), and “budding” tubes forming orthogonally to the epithelial plane, e.g. salivary glands (Chung, Kim, and Andrew 2017; Sanchez-Corrales, Blanchard, and Röper 2018).

As organizing signals such as WNT can induce and orient PCP (Chu and Sokol 2016) within the apical plane, we asked if it is in principle possible to design PCP constraints (not limited to the apical plane) that would result in “wrapping” and “budding”. First, pointing PCP out of the plane was sufficient for a sheet of cells to bend (Figure S11). This is because in our formulation, PCP can drive reorientation of AB polarity and that in its turn is able to bend the sheet (Figure 7). Both “budding” and “wrapping” were qualitatively captured by the model when the axial and radial anisotropy were set by constraining orientation of PCP for cells within a circle (“budding” in sea urchin) and two stripes of cells (mimicking hinge points in neurulation “wrapping”). While CE was needed for proper tube forming in sea urchin example, the tube formed without CE in neurulation (Figure S11).

Thus, the simulations suggest that “wrapping” in neurulation and gastrulation in *Drosophila* (Figure S11) vs. “budding” in sea urchin and organogenesis (Andrew and Ewald 2010; Zegers 2014) may be outcomes of different constraints imposed on PCP. While it is intriguing to speculate that PCP may be oriented out of epithelial plane directly by organizing signals, this may also be an indirect effect of a sequence of intermediate steps. As the organizing signals not only induce and orient PCP but also drive apical constriction (effectively reorienting AB polarity) the PCP may be gradually oriented out of the original plane of epithelium by the following sequence of events: PCP → apical constriction → tilt in AB polarity → tilt in apical plane → PCP out of original epithelial plane. This less precise but simpler interpretation highlights the fact that PCP may drive many of the alternative mechanisms of tubulogenesis and shifts the focus from the differences in mechanisms driving tubulogenesis to the differences in boundary conditions - a set of constraints imposed on cell polarities by neighboring tissue (e.g. notochord in neurulation and organizer in gastrulation).

### Testable predictions

In addition to our conceptual findings, we propose three testable predictions. First, we predict that two potential mechanisms behind the emergence of folds in pancreatic organoids - matrigel resistance and rapid, out-of-equilibrium, cell proliferation - will result in distinct morphologies. Our results suggest that in case of rapid proliferation, the growing structure will develop many shallow folds close to the surface which later tend to deepen. In contrast, external pressure causes fewer but deeper and longer folds (Figure S7). And further, as organoids grow in size, the number of folds will reach an upper limit when under pressure, however, in case of rapid proliferation, the number of folds will keep growing (Figure 5). Visual inspection of published morphologies seems to support the out-of-equilibrium growth (Greggio et al. 2013; Y. Li et al. 2017). To assess if the growth is out-of-equilibrium in 3D organoids, one can quantify the distributions of cell shapes (Cerruti et al. 2013). Our model thus predicts that quantitative counting of folds and measurements of the fold depth and length relative to the size of the growing structure may discriminate between the alternative hypothesis. Quantification of the folds can be done in *in vitro* organoids by either phase or confocal fluorescence microscopy of whole-mount immunostained samples (Greggio et al. 2013; Y. Li et al. 2017). The fold depth and length can be quantified with the same approach as applied to simulated shapes (Methods) in binarized images of the 3D organoid surfaces. As oganoids cannot be cultured without gel supporting 3D growth, it will be necessary to vary both gel stiffness and generation time to uncouple their respective contribution to the folding. The work by (Y. Li et al. 2017) shows that in brain organoids generation time can be both slowed down and speeded up by either genetic manipulation or by adding small molecule inhibitors of pathways regulating cell proliferation. Unfortunately, changing stiffness of matrigel also changes its biochemical composition and may affect cell proliferation and differentiation. One will have to turn to synthetic hydrogels, where it is now possible to uncouple mechanical and biochemical clues (Gjorevski et al. 2016). To illustrate possible applications of our approach, we have only focused on two out of several possible mechanisms that may contribute to folding. The other likely alternative is that folding may result from differences in biomechanical interactions or generation times characteristic to the different cell types. These alternative scenarios are straightforward to consider in our model and will be an exciting venue to explore when more quantitative data on differences in organoid morphologies is available.

Our second prediction is that in case of tubes formed by non-proliferating cells, the length and width of the tubes are controlled by the relative strength of AB polarity and PCP. This result calls for quantification of adhesion proteins along the AB polarity and PCP axes. In PCP mutants with shorter and wider tubes one would expect less planar polarization in adherens junctions and actomyosin, e.g. larger spread compared to wild type in their orientation quantified relative to the tube axis (Nishimura, Honda, and Takeichi 2012). Alternatively, the balance between AB polarity and PCP can be altered by weakening AB polarity, by e.g. mutating tight junction proteins should result in longer tubes. A similar phenotype has already been reported for *Drosophila* tracheal tube (Laprise et al. 2010). With the recent advancements in *in vitro* systems of tubulogenesis, allowing for easy genetic manipulations and more amenable for quantitative imaging, it may in principle be possible to relate the extent of planar anisotropy in PCP mutants and strength of AB adhesion in tight junction mutants with tube length and diameter. The existing coupling between PCP and AB polarity may, however, make it challenging to tweak one polarity at a time.

Our third, and probably most challenging to test, prediction is on the conditions differentiating between tubes forming perpendicular (e.g. sea urchin gastrulation) or parallel (as in *Drosophila* gastrulation or neurulation) to the plane of epithelium. We predict that the outcome will be defined by the orientation of PCP in the invaginating region and the geometry of the boundaries (circular for budding and axial for wrapping) set by e.g. WNT organizing signals. Recent development in imaging localization of PCP complexes in single cells (J. Wu et al. 2013; Chu and Sokol 2016; Minegishi et al. 2017; Habib et al. 2013) allows monitoring localization of PCP complexes, and thus PCP orientation, in individual cells. By placing WNT-soaked (Habib et al. 2013) beads or WNT-secreting cells (Chu and Sokol 2016) one can vary PCP orientation in the cells at the epithelial boundary facing WNT and test for the direction of the epithelial bending and possibly tube formation.

Our approach is by purpose phenomenological and by its nature cannot make predictions about specific molecular details. In all cases, we do not see our simulations as finalized predictions, but rather as pointing in the most promising direction for further exploration of these complex developmental processes. Our setup easily allows for changes as we learn more. The proposed tool should be used in close collaboration with gained experimental knowledge on initial conditions, cell generation times and differentiation processes where polarities play a central role.

Our results open for a series of biological generalizations both in development and diseases. On one hand, we now may be able to explain and unify the apparently very distinct morphological transitions during gastrulation in flies, frogs, fish, mice, and humans by accounting for different initial and boundary conditions. Our model suggests how a moderate change in expression of polarities during some critical evolutionary stages could lead to widely different final morphologies. Thereby, development driven by cell-cell polarity interactions could provide major morphological transitions from local and transient modulations in polarity.

On the other hand, it becomes possible to think of gastrulation, neurulation, tubulogenesis, and organogenesis as the same class of phenomena, where the orientation of the tube is guided by local organizers, and lengths/widths of the tubes are determined by the relative strength of AB polarity and PCP. At the same time, there is an emerging view that wound healing and cancer are local perturbations - e.g. local loss of cells, dysregulation of cell polarities (Martin-Belmonte and Perez-Moreno 2011), proliferation, or autonomously induced organizing signals - of otherwise conceptually the same developmental processes (Humphries and Mlodzik 2017). The power of our model is that it allows to address these hypotheses through predictive models for the dynamics of many cells that interact through combinations of AB polarity and PCP.

## METHODS

Throughout the paper, we use the Euler method to integrate the ordinary differential equations stated in the Model section with dt set to 0.1 or 0.2. Lower dt values will give qualitatively similar results but with increased simulation time. Higher dt values, will result in a collapse of the presented morphologies. The time unit is arbitrary, and the same throughout the paper.

The noise parameter *η* = 10^−4^ where nothing else is stated. Lower noise values will give more smooth simulations, while *η* on the order of 10° will result in collapsing shapes (see also Figure S2E-F).

### Generation time in growing organoids

In Figure 5, the number of cells, *N*, at a given time, *t*, is *N* = 200 exp(ln(2) *t* / *t*_*G*_) where *t*_*G*_ is the generation time. In these simulations, the AB potential (Equation 6) between cells is set to zero when the angle between their polarity is larger than π/2.

### Modeling resistance from matrigel

In Figure 5, we model resistance from the matrigel by imposing a surface force pointing towards the center of mass. The potential of the pressure in the growth medium is given by

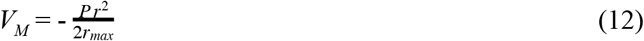

where *P* is the stiffness of the medium, *r* is distance from the center of mass, and *r*_max_ is the distance to the cell that is the furthest away from the center of mass. The resulting force will be constant in time at the periphery. Thus, all cells on a growing sphere will be exposed to a force of equal size. However, cells that end up deep inside a folded morphology will experience weaker resistance.

### Quantification of the local minima

In Figure 5, the number of local minima is defined as the number of cells that do not have any neighbor cells that are closer to the center of mass than themself, and at the same time have an average angle between their AB polarity and their neighbor cells displacement vector that is less than π/2.

### Measuring the tube length and width

In Figure 6, the semi-minor and semi-major axes correspond to the half-width and half-length of the tubes, respectively. As cells on opposite sides of a tube have AB polarity pointing in opposite directions, we approximate the semi-major and semi-minor axes, by finding the half of the maximum and minimum distance between two cells with AB polarity pointing in opposite directions.

### Modeling cells with different polarities

In the gastrulation simulation (Figure 7), each cell is assigned a specific value of polarity strengths (*λ*_1,*i*_, *λ*_2,*i*_, and *λ*_3,*i*_). We define the mutual interaction strength between a pair, *i* and *j* of cells in Equation 4 with different polarity strengths by setting *λ*_1_ = mean(*λ*_1,*i*_, *λ*_1,*j*_) and *λ*_2_ = mean(*λ*_2,*i*_, *λ*_2,*j*_) as well as *λ*_3_ = mean(*λ*_3,*i*_, *λ*_3,*j*_). This choice makes sure that two neighbor cells interact with a force with equal magnitude but opposite sign. Furthermore, it makes sure that *λ*_1_ + *λ*_2_ + *λ*_3_ = 1 holds for all cells.

### Disabling PCP’s effect on AB polarity

In the model described by Equation 6–9, AB polarity and PCP can influence each others orientation. To constrain PCP to the apical plane and thus disable its influence on AB polarity, we set *λ*_2_ = 0 when updating AB polarity in time (during the numerical integration of Equation 10).

### Modeling bottle cells and apical constriction by an external force

The invagination in gastrulation is implemented by adding an external force, *F*, that act on the AB polarity in addition to our usual intrinsic forces from Equation 10

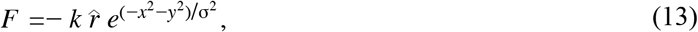

The two parameters are *k* which is the strength of the force (in Figure 7, *k* = 0.02), and σ which defines the decline of the gaussian force (in Figure 7, σ = 10). 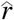 is the unit vector pointing from *x* = *y* = 0 to the cells’ position, and *x* and *y* are the respective coordinates of the cells. This force is applied on the AB polarity and bends the orientation of the polarity away from a *z*-axis (the anterior-posterior axis).

## ACKNOWLEDGEMENT

We would like to thank Anne Grapin-Botton for insightful discussions on organoids and Sigurd Carlsen for discussion on tube formation. This research has received funding from the Danish National Research Foundation (grant number: DNRF116) and the European Research Council under the European Union’s Seventh Framework Programme (FP/2007 2013)/ERC Grant Agreement number 740704.

## SUPPLEMENTARY MATERIAL

### Model details

In our model, we use the following potential to describe the pairwise interaction between cells

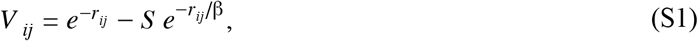

where *r*_*ij*_ is the center-center distance between cell *i* and cell *j*, and *S* is the polarity factor

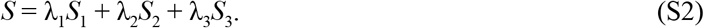

Here, *λ*_1_, *λ*_2_ and *λ*_3_ are the strengths of the respective polarity terms which are given as

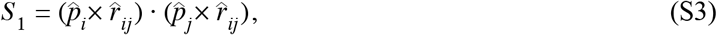

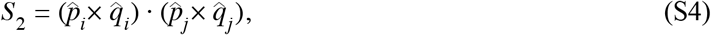

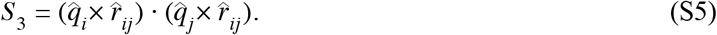

The unit vectors 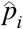, 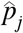 and 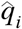 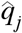 represent the apical-basal polarity and planar cell polarity of cell *i* and *j* Throughout the paper, β is a constant which we set to 5. In order to use the Euler method, we need the gradient of *V*_*ij*_ differentiated with respect to position, 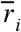, and the two polarities, 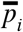 and *q̅*_*i*_:

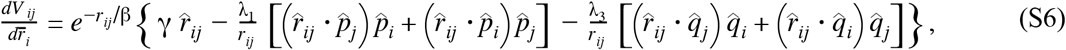

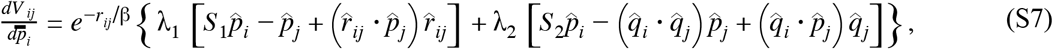

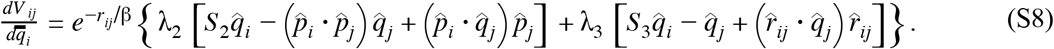

In order to derive Equations S6, S7, and S8, we have used the following:

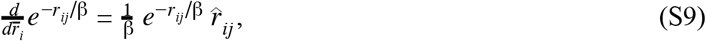

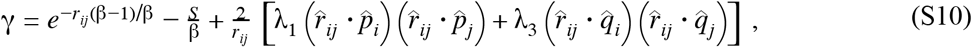

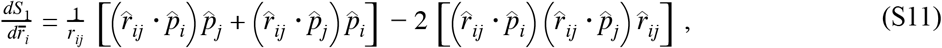

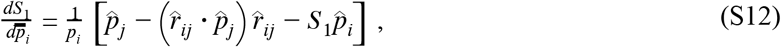

where *p*_*i*_ is the length of the polarity of cell *i* which is equal to one at all times.

**Figure S1.**
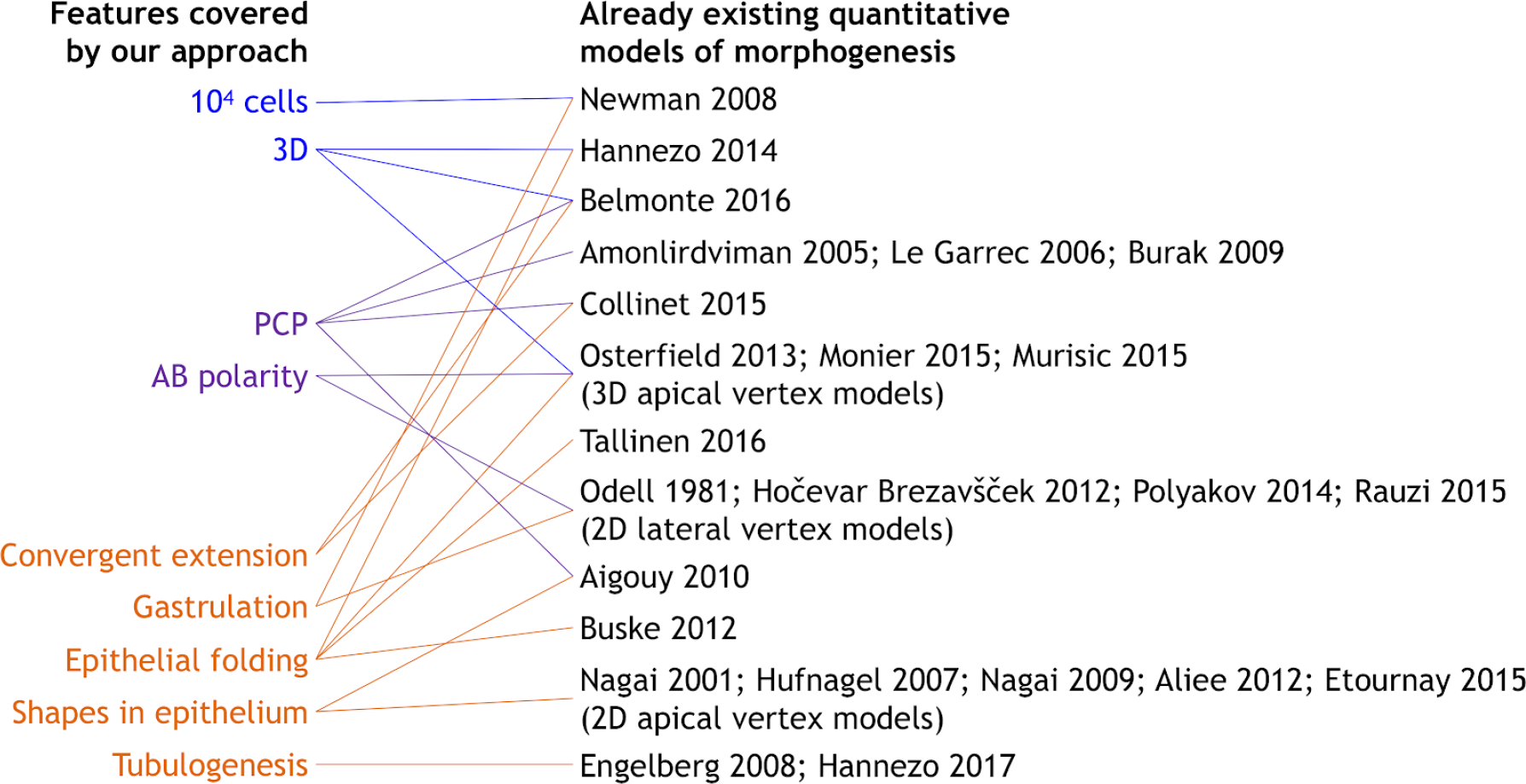
Overview of the existing literature on models addressing specific developmental events discussed in our work. For more references on vertex models see Alt et al. (2017).

**Figure S2.**
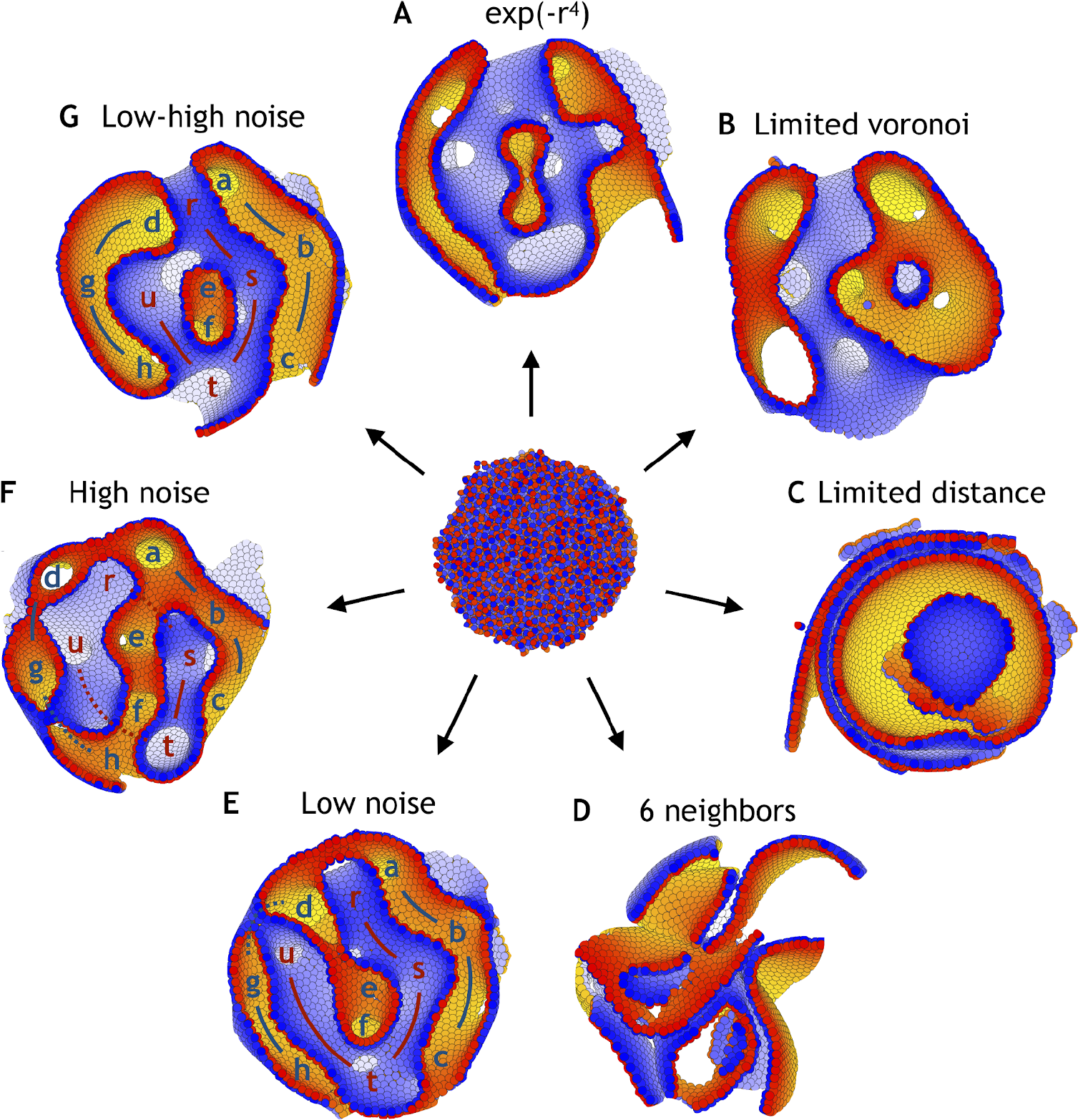
Dependence on the shape of the physical potential, the interaction partners, and noise. **(A)** Applying the neighborhood function shown in Figure 2, but changing the shape of the potential to the short range potential written in Equation 2, the system unfolds and reaches a stable state (*η* = 10^−4^). **(B)** Full Voronoi interactions with a cut-off does also lead to a stable state, although a few cells might lose interaction with the majority of cells (cut-off at 3 shown). **(C)** However, a simple cut-off (and no Voronoi) does not result in stable morphologies but broken sheets on top of each other (cut-off at 2.5 shown). **(D)** The same happens, when all cells always interact with their six nearest neighbors. **(E)** With changed initial conditions compared to Figure 3, the system reaches a different stable state with low noise (*η* = 10^−4^). **(F)** However, this is comparable to the state obtained under high noise (*η* = 10^−1^). **(G)** Increasing the noise in (E) at time log(t) = 4 from *η* = 10^−4^ to *η* = 10^−1^, it results in fewer permutations than having high noise during the entire simulation which is shown in (F). The lower case letters in (E-G) point at macroscopic features (a-h on one side in blue and r-u on the other side in red). The initial positions and polarities are identical for all seven simulations.

**Figure S3.**
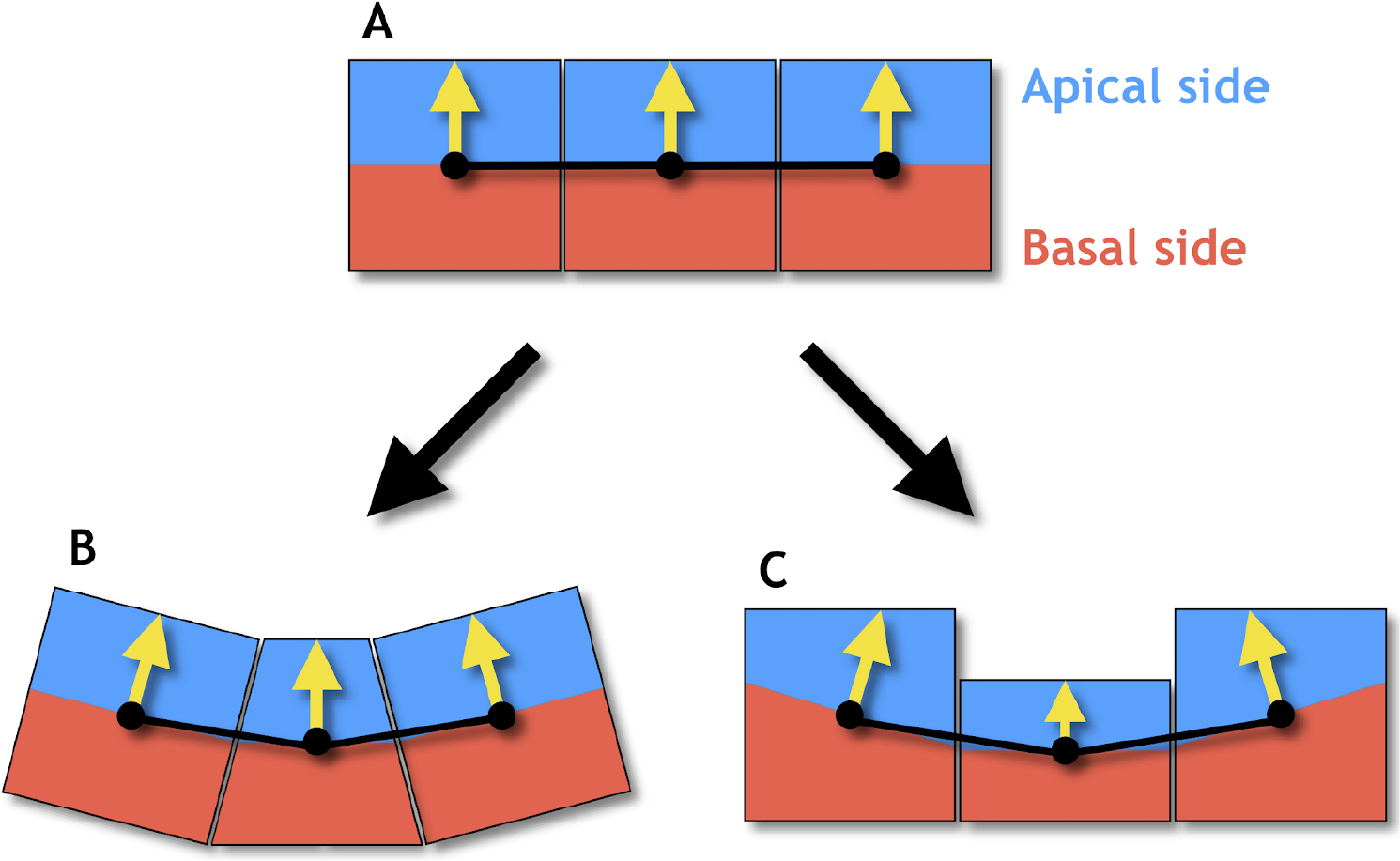
Changes in cell shapes may reorient the orientation of apical-basal (AB) polarity. **(A)** In an epithelial sheet AB polarity (yellow arrows) points perpendicular to the sheet. **(B)** Regulated changes in planar cell polarity can reorient the AB polarities in neighboring cells by apical constriction (narrowing of the apical side) giving rise to a central bottle cell. **(C)** If the AB polarity is shortened, the neighboring cells will reorient in a similar way. These results are obtained under the assumption that the AB polarity tends to orient perpendicular to the distance vectors (black lines) connecting cells’ centers of masses (black dots) which are central to the model presented.

**Figure S4.**
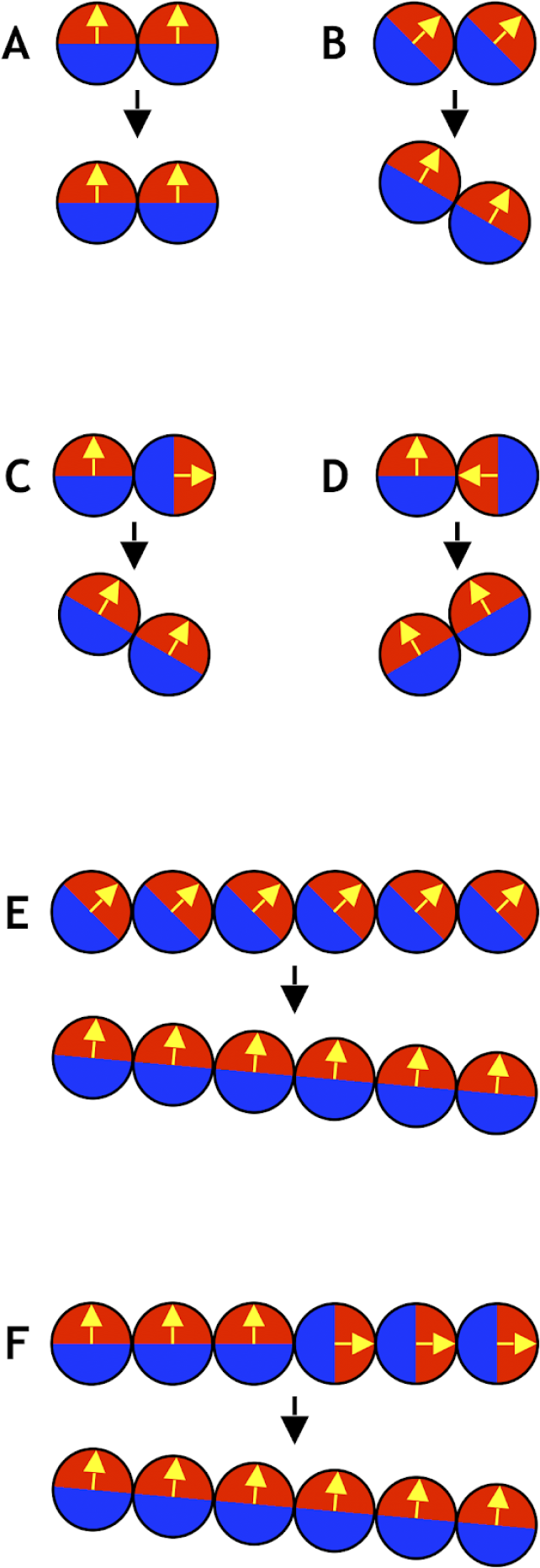
Six simple examples of systems consisting of only two or six cells (see also Movie 1). **(A)** Two cells initially aligned do not result in any movement. **(B)** If both cells’ polarities are 45 degrees to the plane, the axis of position becomes tilted by 30 degrees. **(C)** If one cell points away from another in a two-cell system, it tilts the axis of position. **(D)** Similar to (C), if one cell points towards another, it tilts in the other direction. **(E)** Similar to (B), but with six cells instead of two. In this case the final axis of position is only tilted by 5 degrees compared to the initial axis. **(F)** Similar to (C) and (E), with three cells pointing up and three cells pointing to the right. This also gives a final axis of position that is tilted by 5 degrees compared to the initial axis.

**Figure S5.**
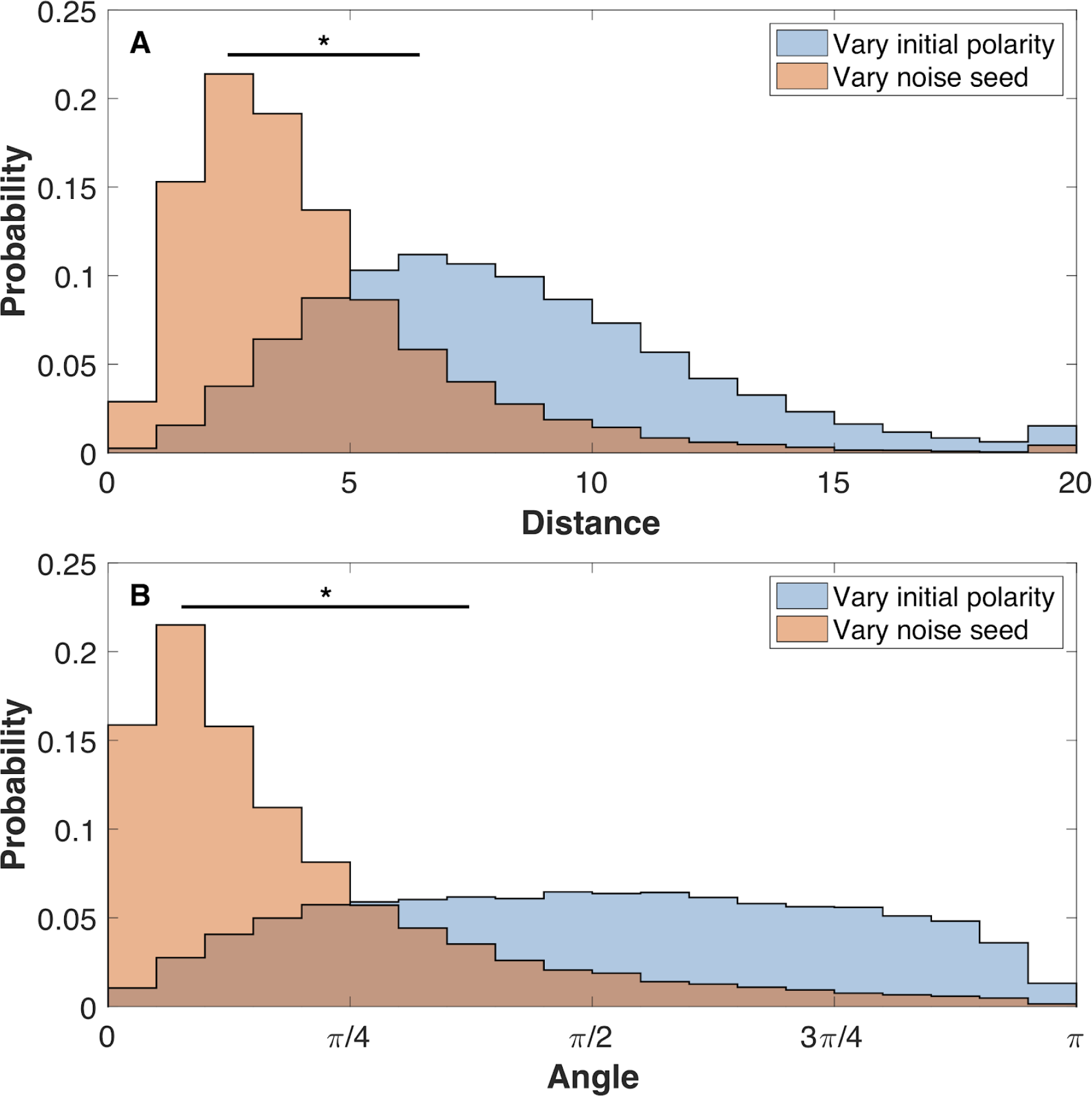
The final shapes are more sensitive to initial polarities than to noise. **(A)** The pairwise distance between cells for three systems with identical initial polarities but different noise and three systems with identical noise but different initial polarities. **(B)** For the same set of aggregates the angle between the pairwise polarities is calculated. The initial positions are the same for all systems. Each system has 8000 cells. Cells are pairs if they were initiated with identical position. Here, the noise level is *η* = 10^−4^. Two-sample Kolmogorov-Smimov tests showed *p* < 0.001 statistical significance (marked by *). Comparing noise levels give similar results as comparing noise seed. The initial polarities are random like in Figure 3.

**Figure S6.**
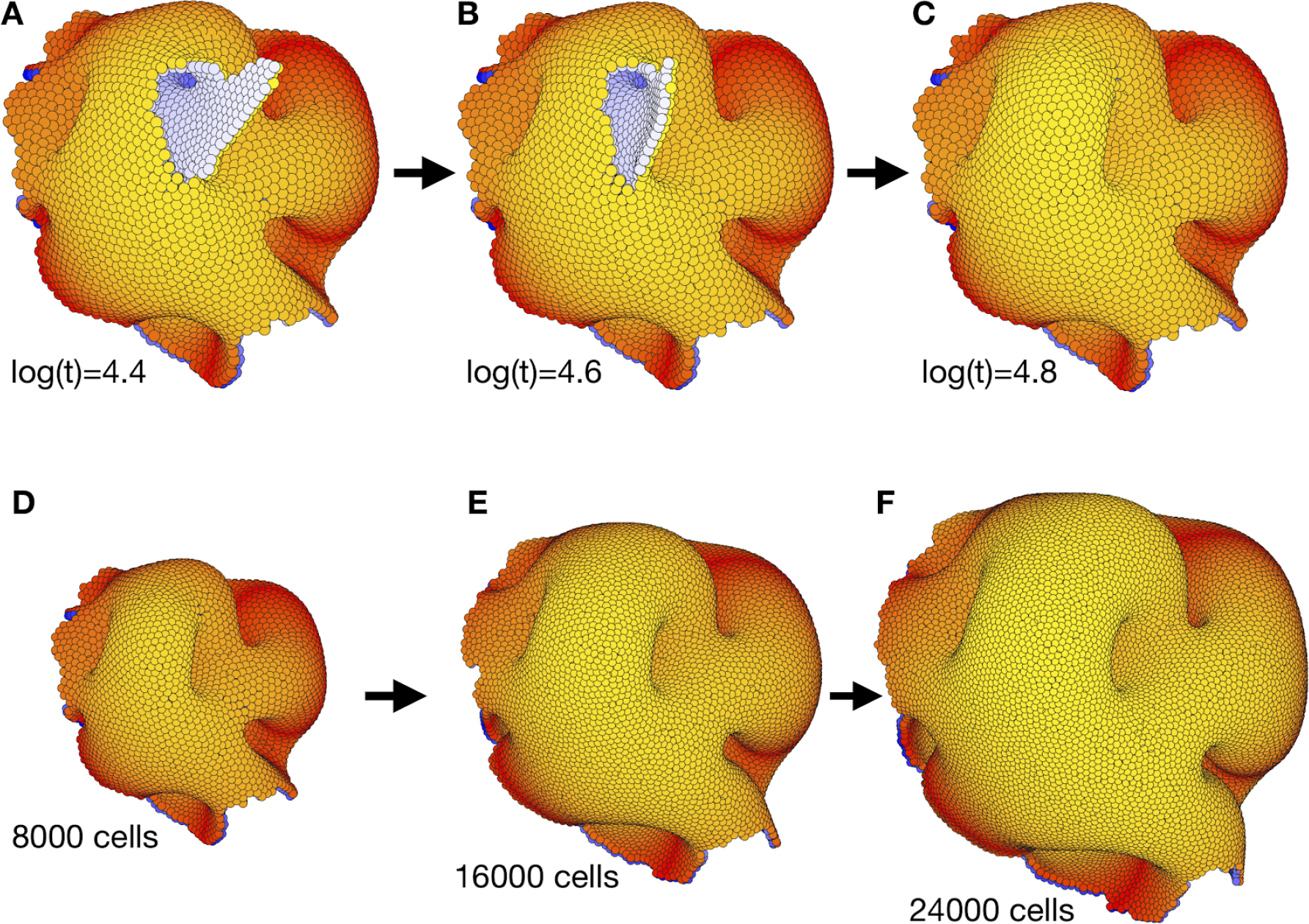
The complex morphology in Figure 3 is robust to puncture and overall system growth. **(A-C)** Self-sealing properties of polarized cell surfaces when close to a final stable state in Figure 3. While the internal morphology remains the same from time log(t) = 3.6 (Figure 3C-D and Movie 2) some of the outer surfaces subsequently reorganize to form a less disrupted torus-like structure with multiple handles. **(D-F)** The final structure in (C) (and Figure 3D-E together with the end of Movie 2) is robust to cell divisions. For every 10th time step, we select a cell by random, and let it divide in an arbitrary direction. We see that the overall shape of the structure is maintained, and that it expands equally in all directions.

**Figure S7.**
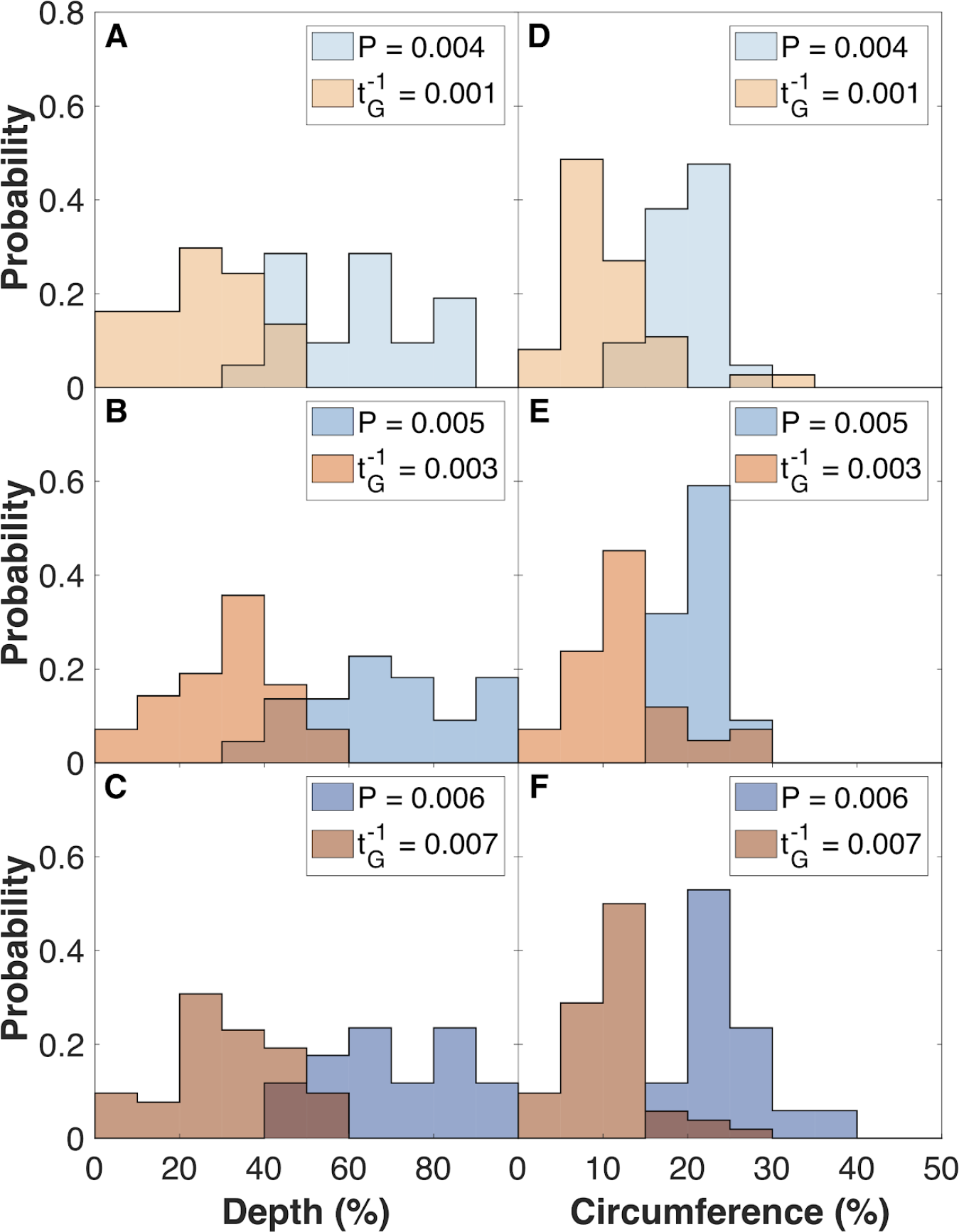
Organoids grown under external pressure have deeper and longer folds compared to organoids grown with rapid cell proliferation. To quantify the folds, we fill the surface of the organoids with water until halfway between the maximum and minimum radius of the system. Then we measure the relative depth and circumference of these ‘lakes’. **(A-C)** Deepest point of the ‘lakes’ (folds) relative to the water level. The probability of having a lake at a given depth is normalized to the number of ‘lakes’. **(D-F)** Length of the ‘lakes’ relative to the entire circumference at this same level. Length of a lake is defined from the angle between the two cells at lake shore that are the furthest away from each other. Pressure and 1/(generation time) increase from upper to lower panels. Two-sample Kolmogorov-Smimov tests showed *p* < 0.001 statistical significance (marked by *). The shown histograms are for the 16.000 cell stage, which compares to the dark blue line in Figure 5.

**Figure S8.**
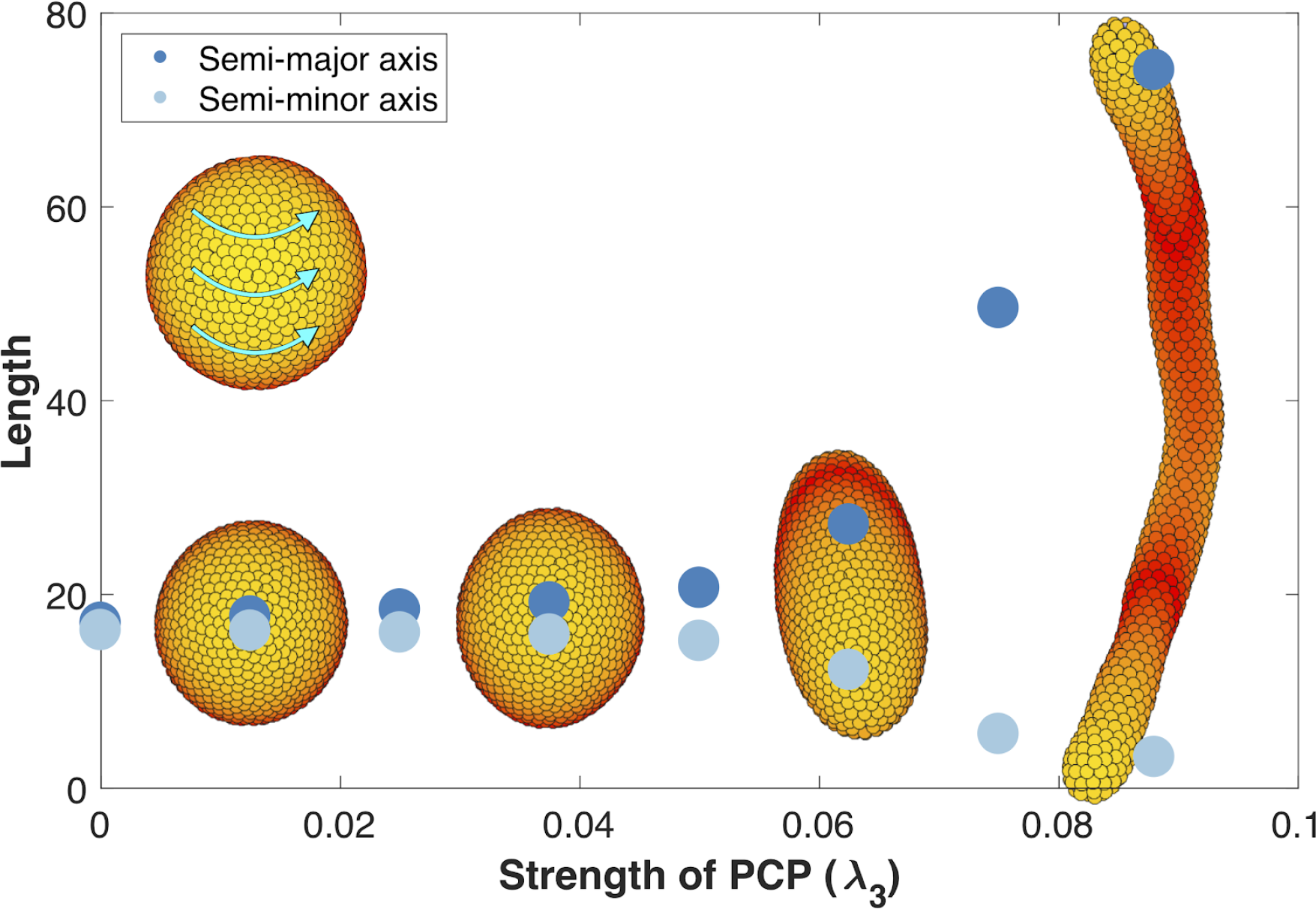
Removing the influence of planar cell polarity (PCP) on apical-basal (AB) polarity. This figure is identical to Figure 6 with the only difference that now *λ*_2_ = 0 when updating AB polarity (*λ*_2_ = 0.5 when updating position and PCP as in Figure 6). The strength of PCP (*λ*_3_) is defined as shown along the x-axis. *λ*_1_ = 1 - *λ*_2_ - *λ*_3_ for updating position and PCP, and for updating AB polarity *λ*_1_ = 0.5 - *λ*_3_. This way *λ*_1_ and *λ*_3_ are the same for position, AB polarity, and PCP, and the only change is the value of *λ*_2_. The final tubes are slightly wider and shorter compared to Figure 6 since the tips become more rounded when PCP does not affect AB polarity. Throughout, all the simulations in this figure, dt = 0.2 and the noise parameter *η* = 5·10^−5^.

**Figure S9.**
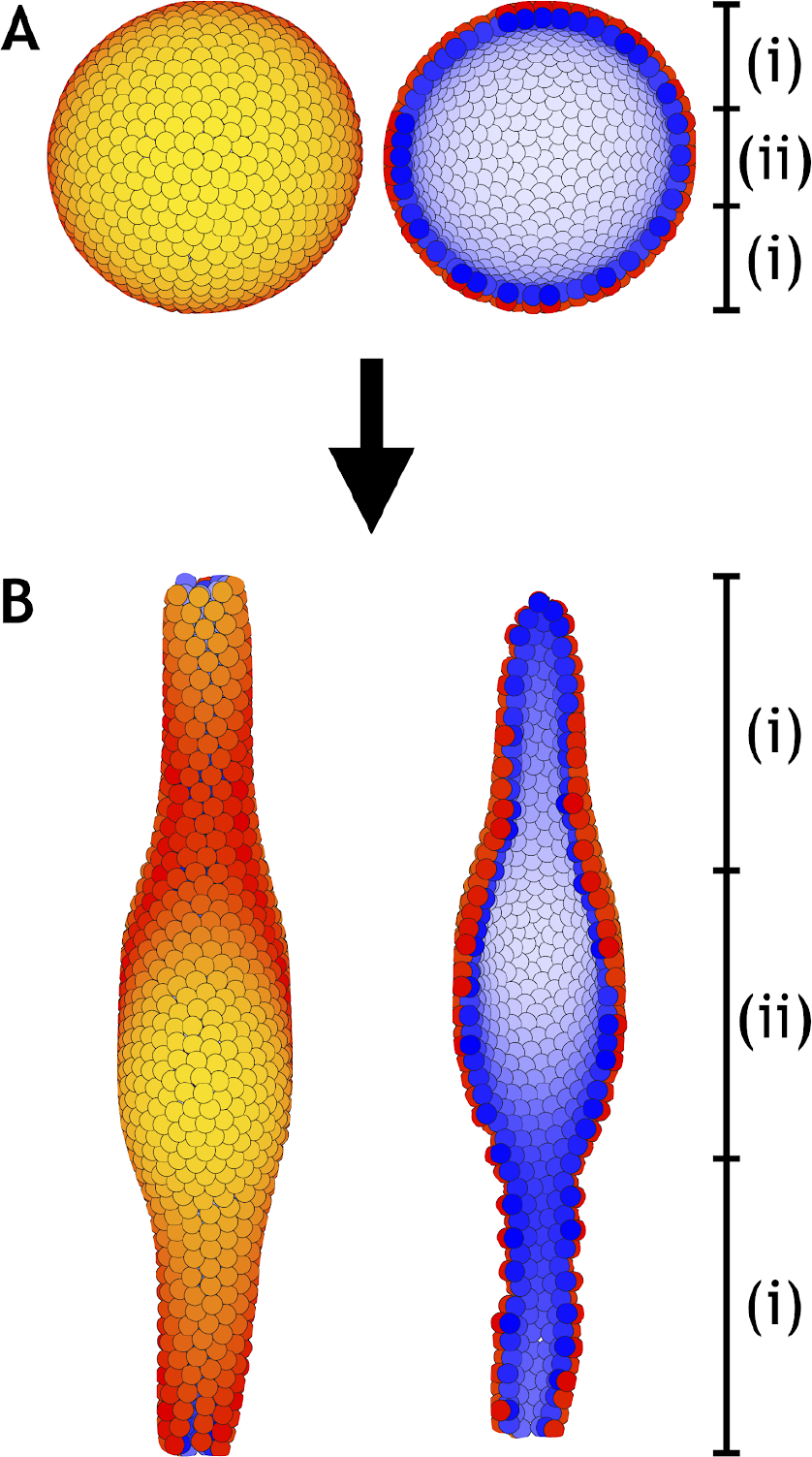
A lumen forms inside a developing tube in areas that lack planar cell polarity (PCP). **(A)** Similar to Figure 6, a hollow sphere of cells is initialized. However, in this example only cells inside zone (i) have PCP while cells inside zone (ii) does not have PCP. **(B)** At the final stage an elongated tube with a central lumen has formed. Images to the left show the entire system while images to the right show a cross section. Cells inside zone (i) develop with *λ*_1_ = 0.41, *λ*_2_ = 0.5, and *λ*_3_ = 0.09 while cells inside zone (ii) develop with *λ*_1_ = 1, and *λ*_2_ = *λ*_3_ = 0. The central third of the 1000 cells in the system belong to zone (ii). Throughout, the simulation dt = 0.1 and *·* = 10^−4^.

**Figure S10.**
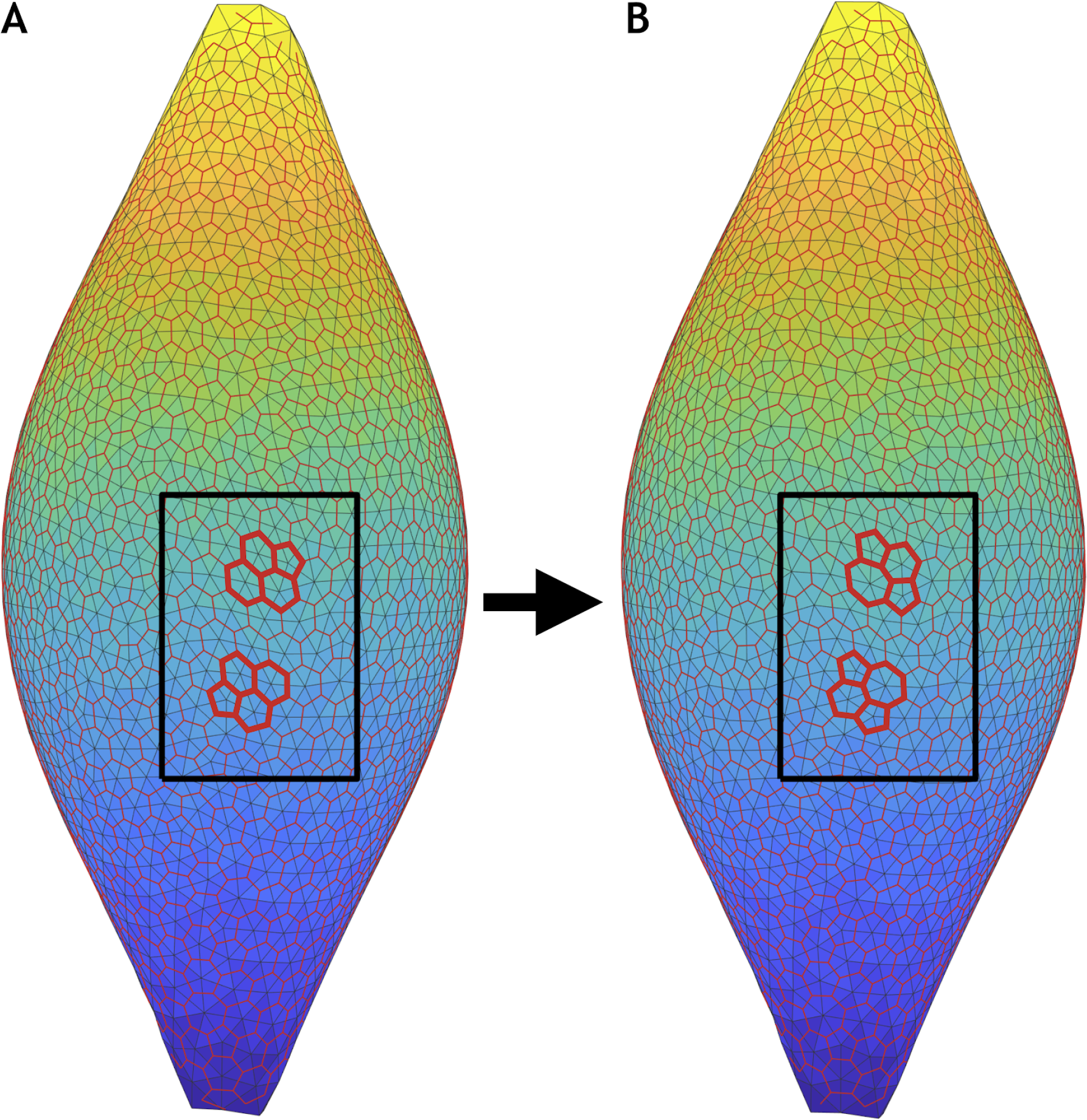
T1 exchanges occur during sphere-tube transition. Two consecutive time frames of the most extreme scenario in Figure 6 (*λ*_1_ = 0.41, *λ*_2_ = 0.5, and *λ*_3_ = 0.09, see also Movie 5). **(A)** Snapshot of the entire system at time *t* = 1259.0. **(B)** Snapshot slightly later at time *t* = 1288.3. In both panels, the cell centers (light grey vertices) are triangulated (light grey edges). Triangle centers (red vertices) are calculated in order to get an approximate location of the cell borders (red edges). Inside the black box, two T1 exchanges (bold red lines) are identified.

**Figure S11.**
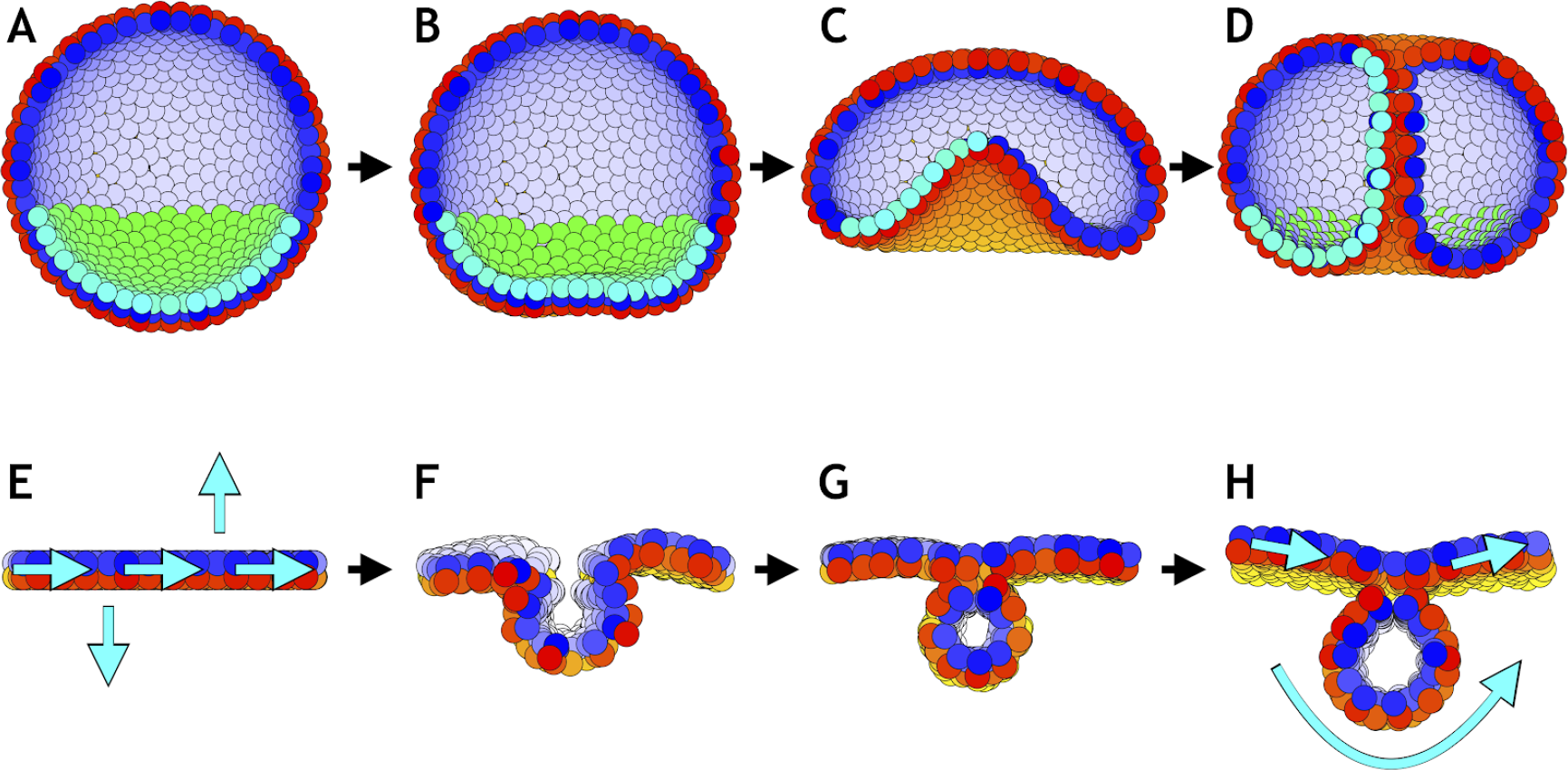
Directed changes in the direction of planar cell polarity (PCP) may drive invagination in gastrulation and neurulation. **(A-D)** Gastrulation in sea urchin modeled without the apical constriction in Figure 7. **(A)** The lower third of the cells in the blastula acquire PCP (cyan-green) pointing opposite to the apical-basal (AB) polarity (red-yellow). **(B)** Flattening of the blastula and invagination occur if direction of PCP is maintained for some time. During this initial phase *λ*_2_ = 0.1, and there is no convergent extension (*λ*_3_ = 0). **(C)** In the final phase, we increase *λ*_2_ to 0.4 and turn on *λ*_3_ = 0.1. With this, we let PCP relax so it curls around the bottom, and allow it to change dynamically in time. **(D)** As a result, tube narrows and elongates, until it finally connects and merges with the top. **(E-H)** Initial conditions on PCP enable neural plate bending and neural tube closure. **(E)** Starting with 1000 cells on a plane with AB polarity, we induce PCP along the plane together with two rows where the PCP points parallel and antiparallel to AB polarity (shown with cyan arrows). Here, the we simulate the neural plate (cells in the middle, between the two rows with constrained PCP) surrounded by the epidermis (the rest of cells). The two rows of cells with PCP pointing out of epithelial plane correspond to the cells at the dorsolateral hinge points next to the neural plate (epidermis boundaries). In chick spinal neural tube can close with only these two hinge points (Nikolopoulou et al. 2017). The bending is driven by apical constriction and PCP is essential for bending, convergent extension and closure. **(F)** This enables neural plate bending and formation of the neural groove. **(G)** Continuing the simulation leads to contact of the two sides of the neural plate and hereby neural tube closure. **(H)** Finally, the system stabilizes with the neural plate on top of the neural tube. Comparing the initial stage to the final stage, the overall direction of PCP in the plate is conserved while in the tube PCP goes around an internal axis. For this simulation, we set *λ*_2_ = 0.5 and *λ*_3_ = 0. Turning on convergent extension (*λ*_3_) at the final stage will allow for elongating the system along the axis going through the tube and narrowing it in another direction. The concept is similar to gastrulation in *Drosophila*. In both simulations, sea urchin and neurulation, dt = 0.3. In sea urchin (A-D), the noise parameter *η* = 3.3·10^−5^, and in neurulation (E-H), *η* = 3.3·10^−2^.

### Movie 1 (Related to Figure S4)

Dynamics for two or six interacting cells without noise (*η* = 0 and dt = 0.1, see also Figure S4). **(A)** No movement occurs if the polarities are perfectly aligned and the distance between the cells is at steady state. **(B)** If both cells initially have their polarities 45 degrees to the axis of position, then both cells rotate slightly, and at the final stage the axis of position has been rotated by 30 degrees. **(C)** Initializing one cell with polarity up and another cell with polarity to the right results in substantial rotation of the latter cell and less rotation of the first cell. Finally, the cells reach steady state positioning each other on a tilted axis. **(D)** Opposite (C). One cell has polarity up while another cell points left. This results in an axis that is tilted to the left. **(E)** Similar to (B), but with six cells having their polarities initially 45 degrees to the axis of position. In this case the outermost cells move the most and rotate slower than the four inner cells. Finally, resulting in an only slightly tilted plane. **(F)** Similar to (C) and (E). On the left three cells point initially up while on the right three cells point initially to the right. In this case the three cells on the left almost do not rotate and move at all while the three cells on the right rotate and move substantially.

### Movie 2 (Related to Figure 3)

An aggregate of 8000 cells all with apical-basal polarities initially pointing in random directions develops into a final stable complex morphology. During the simulation the polarities and positions are updated dynamically with equal speed and noise (dt = 0.1 and *η* = 10^−3^). There is no planar cell polarity (*λ*_1_ = 1 and *λ*_2_ = *λ*_3_ = 0). **(A)** The entire system unfolds, and we notice that the outer surface reaches equilibrium state later than the internal morphology. **(B)** Cross section of the system at *y* = 0. **(C)** Same as (B) but viewed from an angle slightly above and rom a side. The color scheme is as described in Figure 3.

### Movie 3 (Related to Figure 4)

Dynamics when the polarities have restricted orientations. In all three simulations, an aggregate consisting of 8000 cells develops into three simple morphologies due to polarities being fixed in different directions. **(A)** One big lumen forms when the polarities are restricted to point radially out from the center-of-mass. At an early stage, a small lumen forms around the center-of-mass. Later the shell transiently consists of multiple cell layers. However, in the end these layers merge into a single cell layer (Figure 4A). **(B)** A tube emerges when the polarities are restricted to point away from a central axis. Similar to (A), a small tube appears at an early stage deep inside the aggregate, and transiently it contains multiple cell layers (Figure 4B). **(C)** Finally, two opposing planes form when the polarities are restricted to orient away from a central plane. This develops in a similar way as (A) and (B) without rotating any cells (Figure 4C). Throughout all three simulations, dt = 0.1 and the noise parameter *η* = 0.1.

### Movie 4 (Related to Figure 5)

*In silico* organoids grown from 200 up to 16000 cells. **(A)** Increasingly many surface-near folds emerge when the organoid is grown with rapid cell proliferation (*t*_*G*_^−1^ = 7·10^−3^, *P* = 0. Figure 5A). The image at the top illustrate the entire system, while the image at the bottom shows a cross section of the system. **(B)** A saturing number of deep folds emerge when the organoid is grown slowly in a resistant matrigel (*t*_*G*_^−1^ = 1.4·10^−4^, *P* = 0.006, Figure 5B). As in (A), the entire system is presented on top of a cross section of the system. Throughout, all organoid simulations in Figure 5 including the ones shown in this movie, *·* = 10^−4^ and dt is the minimum of 0.2 and one fifth of the time to the next consecutive division in the system.

### Movie 5 (Related to Figure 6)

Model of tubulogenesis. A spherical lumen consisting of 1000 cells with apical-basal polarity pointing radially out gets planar cell polarity (PCP). In equilibrium, PCP will curl around an internal axis. Depending on the relative strength of the polarities the sphere will elongate and transform to a tube with a given length and width. This movie illustrates the most extreme scenario in Figure 6 (*λ*_1_ = 0.41, *λ*_2_ = 0.5, and *λ*_3_ = 0.09). The left panel shows the entire system while the right panel shows a cross section of the system. Throughout all simulations in Figure 6 including the one presented in this movie, dt = 0.2 and *η* = 5·10^−5^. The color scheme is as described in Figure 3.

### Movie 6 (Related to Figure 7)

Model of sea urchin gastrulation. Starting from a hollow sphere of 1000 cells with apical-basal polarity pointing radially out. The bottom flattens and invaginates by applying an external force to mimic apical constriction (see Methods), and by giving a fraction of cells planar cell polarity pointing around the sphere. The tube forms, elongates, and connects with the upper part of the blastula. Throughout the simulation, dt = 0.1 and *η* = 10^−4^. For more details see the caption to Figure 7. Panels are similar to Movie 5. The color scheme is as described in Figure 3 and Figure 7.

### MatLab script

MatLab script to generate and visualize data. MatLab R2016b or newer is required together with the Statistics and Machine Learning Toolbox. In addition, the Parallel Computing Toolbox is required if the *PAR* parameter in the basic script is set to 1. The input folder contains initial conditions for three standardized systems. Bulk systems have neither apical-basal (AB) polarity nor planar cell polarity. Plane and shell systems have only AB polarity. ‘N’ in the file names gives the number of cells in the system. The initial polarity directions can be modified on line 4-5 in the basic.m file. Inside this file, it is also possible to set the degree of noise (*η*). the size of the time steps (dt), and the relation between the polarity strengths (*λ*_1_, *λ*_2_, and *λ*_3_ The parameter *inc* is used to speed up the simulations by only applying the neighborhood function to the nearest 100 neighbors. Generated data is saved in the output folder, and the visualization script is in a seperate folder.

